# Evaluation of analysis modes for RNA coexpression in single-cell and bulk tissue

**DOI:** 10.64898/2026.06.17.731965

**Authors:** Ching Pan Chu, Nairuz Elazzabi, Jules Garreau, Brianna Xu, Xinrui Xiang Yu, Alexander Morin, Paul Pavlidis

## Abstract

Coexpression of transcripts presents the most common means of computational inference of transcription factor regulation, and is often combined with other data types to infer regulatory networks. With the growing popularity of single-cell approaches, there are questions about how best to extract coexpression information from the data. Recently we reported a simulation study that explored the differences among coexpression performed at different levels: across single cells (xCell, per cell type), across subjects from pseudobulked single-cell data (xSubject, per cell type), or across subjects using bulk tissue samples (xBulk). Here we test predictions made by those models using real data. We consider both preservation (consistency of coexpression findings across different levels of analysis of the same data) and replicability across independent studies, as well as biological interpretability. We find that preservation across levels is limited, indicating the choice of analysis level will affect outcomes. We show that xCell coexpression is more replicable across studies compared to xSubject. xBulk coexpression is dominated by patterns driven by variability in cellular composition and fails to capture much coexpression that is reliably detected at finer resolutions. While all modes of analysis exhibit some enrichment for known regulatory relationships, it was highest with the xCell mode. Finally, we present a case study of the effect of analysis modes on a schizophrenia-associated pattern, reinforcing the importance of analytic choices in the interpretation and replicability of coexpression analyses. Together with our modeling study, this work emphasizes the importance of understanding sources of expression covariation as they relate to the goals of the analysis, and recommend single-cell-based data with biological replicates should be the focus of attempts to infer dynamic regulatory interactions that are more likely to be replicable by others.

## Introduction

One of the main goals of genomics is to decipher regulatory circuitry that underlies gene expression. A bedrock method is the use of RNA coexpression analysis, based on two ideas: First, that genes which are co-regulated will tend to show correlated transcript levels; and second, correlations between the transcript levels of a transcription factor (TF) and another gene is a piece of evidence that the gene is a target of that TF. The latter mode has particularly gained popularity. Methods like WGCNA (Langfelder and Horvath, 2008) encourage the identification of “hub” genes that are often interpreted as key drivers of the observed patterns, while methods such as SCENIC (Aibar et al., 2017) look directly for coexpression between TFs and potential target genes. It is also well known that coexpression alone is insufficient to infer a direct regulatory relationship, because of its correlative nature. Data such as TF motifs near a putative target gene are commonly employed as independent evidence and can be used to infer a directionality of the regulation. While there is evidence from formal assessments that such methods don’t have very high specificity or sensitivity (Marbach et al., 2010; Pratapa et al., 2020), it is still common for researchers to build GRNs and analyze and interpret them with confidence, and some success. Improving the quality and interpretability of coexpression analyses would be beneficial.

In our recent work (Chu et al., 2025) and in this paper, we are examining the foundations of regulatory inference from coexpression. We started by considering coexpression as emerging from regulation happening within individual cells. Ideally, one would be able to repeatedly sample a cell and directly detect coexpression over time, due to changes in state driven by internal or external signals, as well as stochastic effects that still reflect the underlying regulatory network in that cell. We refer to this type of regulation as “dynamic” to differentiate it from more static differences such as between two disparate cell types (while acknowledging this distinction may not always be clean).

Because transcriptome-wide measurement over time in single living cells is not yet technically feasible, cells are destructively sampled, yielding a snapshot of the transcriptional state (“pseudotime” methods can be used to try to infer temporal information from a single cross-sectional sample, but we don’t consider this type of experimental design further here). In our conceptualization, which we believe aligns with the intentions of most researchers using coexpression, the primary interest is dynamic regulation due to events occurring within cells.

Researchers have multiple ways to study coexpression, given a starting point of an experimental design sampling from multiple subjects (Figure 1; we use the term “subject” to indicate an entity such as an individual human or other organism that was sampled). We are referring to differences in how the data are collected and organized to produce the input expression matrix. These choices result in different units of analysis, and thus define one dimension of the expression matrix (the other being defined by the genes). First, one can use bulk tissue samples (xBulk), where each sample typically comes from a different experimental subject. This has been done in many thousands of publications and continues to be very popular. For example, the WGCNA R package (Langfelde and Horvath, 2008) is used almost exclusively for bulk data analysis and was cited over 4500 times in 2025 alone based on Google Scholar citation metrics.

**Figure 1:**
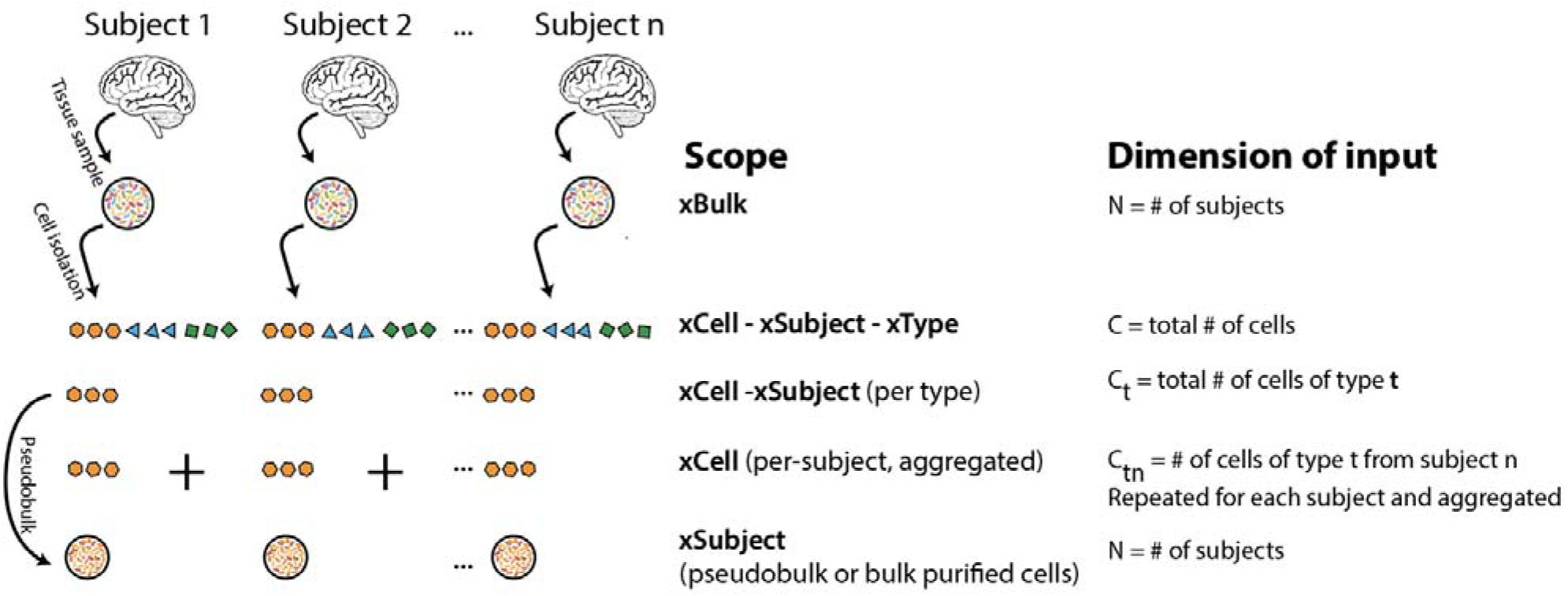
Modes of coexpression analysis. The schematic at left shows the steps and data inclusion for each of the modes listed at the right. Cell types are indicated by colors and shapes; data from each subject is organized from left to right. Bulk samples or pseudobulks are indicated by larger circles. The dimensionality of the input for coexpression analysis is described, referring to the columns of the gene x sample expression matrix.

Alternatively, and increasingly, researchers can use single-cell (or single-nucleus; henceforth scRNA-seq or single-cell) methods. In contrast to bulk samples, scRNA-seq affords multiple options. The obvious one, and perhaps the most common, is to consider all the cells at once as individual experimental units. We refer to this approach as xCell-xSubject-xType, because the expression matrix contains all the cells as columns regardless of subject and type/states (we use the term cell type for simplicity). This is the standard approach for SCENIC. A variant is xCell-xSubject, where the SCENIC-style approach is applied separately to each cell type. While this appears uncommon, it was used in at least one publication (Ling et al., 2024). A second major option is to pseudobulk each sample, per cell type, yielding multiple expression matrices where cell-level information has been lost. We call this xSubject, because the unit of analysis is the subject. This approach is also less commonly used, with a prominent example also being the work of Ling et al (2024). It is possible to consider combining all cell types in this approach (xSubject-xType) but we are not aware of this being applied so do not consider it further here. The third approach, xCell, is simple if one has a single subject: for a given cell type, the columns of the expression matrix are the individual cells. When there are multiple samples, we construct these matrices for each subject (for that cell type) and aggregate the resulting coexpression matrices, in effect taking an average of the coexpression found in each subject (see Supplementary Figure 2 for a schematic). This explicitly discards covariation that might exist across subjects. The xCell method (or variations of it) have been used in several publications, including our own work (Harris et al., 2021; Morin et al., 2025). Further explanations of the xCell, xSubject and xBulk modes can be found in Chu et al. (2025) and other details are in the Methods section.

In this work, we focus on comparing three of these modes: xBulk, xCell, and xSubject. We examine several hypotheses. Our first is that the coexpression signals one detects are not equivalent across the modes, and in particular, signals detected at the xCell level will often not be identified at the other levels of analysis. We refer to this as *preservation* (or lack thereof). Our second hypothesis is that *replicability* across studies will also be different. Specifically, we predict that xCell coexpression analysis is more replicable across independent studies, because approaches that incorporate xSubject sources of transcript covariation might yield results that are idiosyncratic to the subjects used in the study. We further hypothesize that the biological interpretation of coexpression will not be the same, in terms of predicting TF-target relationships.

In our modeling study (Chu et al., 2025), we explored the preservation hypothesis with simulations using empirically-derived parameters. We showed that it is unlikely that coexpression patterns present at the cell level (xCell) will directly survive to be detectable in single cell coexpression analysis across subjects (xSubject) or in bulk tissue (xBulk), unless very particular conditions are met. However, it is unclear the extent that such conditions are fulfilled in practice. For example, a dynamic regulatory mechanism that underlies xCell coexpression could in theory give rise to covariation across subjects. In the extreme possibility that coexpression patterns at all different levels of resolution and among different cell types are concordant and synchronized, xCell coexpression signals would be highly preserved. It has been reported that there is some concordance between xCell and xBulk coexpression (Farahbod and Pavlidis, 2020; Harris et al., 2021). Since bulk tissue data is generally easier and less expensive to gather than single-cell, their equivalence would raise the question of the value of single-cell data for coexpression analysis. These studies did not explore xSubject variability within the same cell types, nor the impact of dilution on cell type specific signals in bulk tissue. As such, the relationship between xCell and xBulk coexpression and the reasons underlying signal concordance (preservation) can still be clarified, as well as the issue of cross-study replicability.

Here we compare coexpression at the different levels of resolution and quantify the concordance among them, and examine their ability to recover experimentally validated regulatory interactions. We further use the data of Ling et al. (2024) to present a concrete case study of the impact coexpression analysis decisions have on data interpretation. Together with Chu et al. (2025), this work lays out a framework for understanding coexpression data in its various forms, and further supports the effectiveness of analysis at the cell level (per cell type; xCell) in highly replicated designs as optimal for applications of coexpression to investigating dynamic regulation.

## Methods

Code and data underlying the analyses will be made available upon publication.

### Data preparation

We sought single cell transcriptomics data with replication as well as those with matched bulk tissue samples. We identified seven single cell human brain datasets that met the criteria for inclusion (Supplementary Table 1). Two of these datasets (ROSMAP and Velmeshev) also had matched bulk tissue samples for at least some samples. For all the single cell datasets except the ROSMAP and Velmeshev datasets, we obtained the expression matrices and the corresponding metadata from Gene Expression Omnibus (GEO) using the accessions provided in the original publications. For the Velmeshev dataset, we downloaded the single cell data from the UCSC cell browser website (Speir et al., 2021) (https://cells.ucsc.edu/?ds=autism) on April 7, 2023 and the bulk tissue data were provided to us directly by the authors. We downloaded the ROSMAP single cell and bulk datasets from Synapse (https://www.synapse.org/#!Synapse:syn18485175) on April 7, 2023. All the datasets had cell type annotations. We mapped these to six cell types in the brain: excitatory neurons, inhibitory neurons, astrocytes, oligodendrocytes, microglia, and oligodendrocyte progenitor cells (OPCs).

We refer to the cells belonging to each cell type and dataset as a “data group”, denoted with the notation Dataset:Cell_type e.g Velmeshev:Inhibitory. We preprocessed the cells for each data group separately. For each group, we retained genes expressed in more than 5% of all cells. We also filtered out cells expressing too many or too few genes (top and bottom 5%). The number of genes and cells per data group after filtering are shown in Supplementary Figure 1A and B. Next, within each data group, we filtered out subjects with less than twenty cells, and subsequently data groups with less than twenty subjects remaining. After filtering, all but two datasets retained all six cell types. Specifically, the Lau dataset did not have OPCs whereas the Ramos dataset did not have oligodendrocytes or astrocytes. In the end we retained expression data for 39 data groups. Finally, we normalized all expression values into counts per million reads (CPM). After filtering, we retained a total of 882,527 cells. There was substantial overlap of genes detected in the same cell types among the different datasets and they were also highly correlated in mean expression within each cell type (Supplementary Figure 1C). These checks are consistent with the data sets being generally comparable. Overall, the corpus is of 39 data groups for analysis (7 data sets, 4-6 cell types each).

We processed the bulk tissue data similarly. The ROSMAP bulk dataset had 636 bulk tissue samples, of which 32 samples had matched single cell data. The Velmeshev dataset had 41 bulk tissue samples, all of which had matched single cell data. After quality control, the two datasets retained 21,518 and 25,326 genes, respectively. The genes detected in the two datasets largely overlapped (21,239) and the expression levels across genes were highly correlated (mean Pearson’s correlation r = 0.9, Supplementary Figure 1D).

### Construction of xCell, xSubject, and xBulk coexpression

We computed a xCell coexpression for each data group by first computing a separate coexpression matrix for each subject and then aggregating as described previously (Morin et al. 2025; Chu et al. 2025). Within a data group, the expression profile of gene *A* in subject *s* can be represented as a vector *Exp_cet_s_(A)* of size *C_s_* where *C_s_* is the number of cells from subject *s*. Thus, the pairwise xCell coexpression between genes A and B within subject *s* can be computed as Pearson’s correlation of the vectors *Exp_cet_s_(A)* and *Exp_cet_s_(B)*. Repeating this calculation for all possible pairs of genes within all subjects produces a set of coexpression matrices where each coexpression matrix *Coex_xCel_s_* is a symmetric (g rows and columns where g is the number of genes). Each element in *Coex_xCel_s_* contains the xCell correlation of expressions between the two indexed genes. Finally, we rank normalized the coexpression values by taking the *rank/max(rank)* to enforce a consistent uniform distribution across all subjects. This results in coexpression values that range from 0.0 being the lowest coexpression, to 1.0 being the highest. Importantly, for undetected genes, their coexpression with all other genes were imputed to the median coexpression value, effectively placing them in the middle of ranking.

We computed xCell coexpression for each subject within a given data group, and then aggregated the subject level coexpression by averaging, thereby producing a final matrix for each data group. We performed a final round of rank normalization so that the coexpression values are comparable across data sets.

We computed xSubject coexpression by first collapsing the cells belonging to each subject into a single expression vector (pseudobulking). The subject level expression of A is the mean value across all cells belonging to subject *s*: *Mean(Exp_cet_s_(A))*. Repeating this calculation for all S subjects produces a subject level expression vector, *Exp_sbj(A)*, of size S. Then, we compute the pairwise xCell coexpression between genes A and B across subjects as the correlation of the vectors *Exp_sbj(A)* and *Exp_sbj(B)*. Again, we repeated this calculation for all 39 data groups. In this way, the single cell-level expression profiles were used as estimates of subject-level expression profiles, rather than to compute coexpression directly. Importantly, the xSubject coexpression was computed using the same single cell datasets that were used to compute xCell coexpression. Thus, there was a xSubject analysis for every xCell that used the same underlying data.

For xBulk, we computed xBulk coexpression for each pair of genes, using all of the available samples. For consistency with the xCell and xSubject, we used rank normalized coexpression values at the xBulk level.

### Coexpression replicability and preservation

We used odds ratios (OR) to quantify the extent of concordance among coexpression data sets. This measures the extent of overlap of coexpression edges, which requires binarization of coexpression measures into positive and negative edges. We designated the top-ranked 1% of the edges in each as coexpression edges and the remaining 99% as negatives. Each comparison involves one reference and one test, and we computed an odds ratio that reflects the level the test is enriched for the coexpression edges identified by the reference. Specifically, suppose that the set of positive edges in the reference is denoted *P_ref_*, and the positive set in the test is denoted *P_test_*. Accordingly, the negative edges in the reference and test are respectively denoted *N_ref_* and *N_test_*. Then, odds ratio is given by:

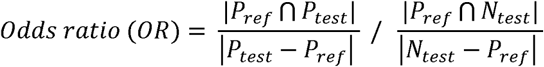

Finally, we computed p-values using Fisher’s exact test to measure the probability of observing the given odds ratio under the null hypothesis that OR = 1. We used odds ratio to measure both replicability and preservation of coexpression signals. For replicability, we compared the coexpression of different datasets to each other, within the same cell type and level of cellular resolution. Since the odds ratio is asymmetric, we calculated an odds ratio for every pairwise permutation of datasets. This means that the replicability between the coexpression in any two datasets was computed twice, with alternating reference and test designations. The mean replicability of a given dataset was the average across all the comparisons where it is used as the reference.

To measure preservation, we compared the capacity for analysis at the lower cellular resolution to recover the edges identified at the higher (finer-grained) resolution. Therefore, in the comparison between xCell and xSubject or xBulk, the xCell coexpression was used as the reference in computing odds ratios. Similarly, for the comparison between xSubject and xBulk, xSubject is used as the reference.

### Comparison with literature curated co-regulatory pairs

To evaluate whether coexpression contained signals of intracellular regulation, we used the catalogue of experimentally validated direct transcriptional regulatory interactions (DTRIs) that we previously assembled [Chu et al.] and have since expanded. DTRIs are TF-target pairs where the TF has been shown to regulate the target gene by low-throughput experimental evidence. Collectively, there are 4,387 annotated DTRIs. Because our curation included annotations of the tissue context, we filtered for those with experimental evidence in the human or mouse central nervous system to focus on relations most likely to be relevant to the tissue used in our experiments. We also filtered out TFs with only one target gene because such records cannot be used to derive co-regulation of targets. After filtering and mapping to Ensembl identifiers, we retained a final set of 247 DTRIs involving 43 TFs and 184 target genes. From this, we derived set of co-regulated pairs of genes, by generating all pairwise combinations of target genes for each TF. The collection of co-regulated gene pairs constituted the positive set. To control for gene expression biases, we designated all other possible pairwise combinations among the same set of genes as the negative set. In total, there were 1,480 gene pairs in the positive set and 15,356 gene pairs in the negative set, involving 184 genes. To measure enrichment, we vectorized each coexpression matrix into a list of gene pairs, ranked by coexpression values. We used the Mann-Whitney U test to quantify the level of enrichment of the co-regulated pairs at the top of each coexpression ranking. Note that because the sets of detected genes are variable across studies, we further filtered the positive and negative sets for each comparison accordingly. Finally, to control for node degree of the co-regulated genes, we generated a null distribution for each comparison by randomly shuffling connections among the co-regulated pairs, while maintaining their node degrees. This was to ensure that we were prioritizing the co-regulated pairs specifically rather than indiscriminately prioritizing highly connected genes. By comparing the observed AUROCs to the null distribution of 1,000 AUROCs, we obtained empirical p-values as an additional measure of confidence.

### SNAP pattern analysis

We obtained the data for Ling et al. (2024) from the authors. This included cell-level counts, cell type annotations and some sample meta-data. We excluded 11 outlier samples as identified by Ling et al., for a total of 180 samples.

Analysis of coexpression of SNAP genes: We considered 643 of the top 1000 SNAP genes reported by Ling et al. that were protein-coding, had expression in our other data sets, and which were reported to be expressed by Ling *et al*. in excitatory neurons and astrocytes. These 643 genes were stratified by the sign of their L4 loadings reported by Ling et al., resulting in 183 genes with positive loadings and 460 with negative loadings. Using the cell-type expression annotations reported by Ling et al., we then combined loadings sign and cell-type expression to define four gene sets: (ast_pos, ast_neg, exc_pos, ext_neg.

We computed the correlation ranks for the four gene sets under each analysis mode (xCell, xCellxSubject, xSubject), in each cell type (astrocytes and excitatory neurons), and compared correlations across the three analysis methods for each gene set and cell type, computing Z-scores for each edge as follows. For each combination of cell type and coexpression analysis mode, we generated a cell type- and analysis mode-specific random background distribution by sampling 1,000 unique edges (without replacement) from the correlation matrix. We then computed Z-corrected correlations for each combination of gene set and mode of analysis and cell type using the appropriate matched background: Z = (correlation - mean_background) / SD_background. To compute odds ratios, we used genes in the peaks of coexpression for the SNAP genes (defined as Z-score > 1; "peak-driving" edges) rather than the top 1% coexpression threshold used in our other analyses. We did this because SNAP genes are not well captured by our top 1% criteria. For instance, only 2.98% of SNAP genes in astrocytes and 0.82% in excitatory neurons had a correlation ranking in the top 1% (at the level of coexpressed gene pairs the overlap is much lower, with Jaccard indices of 0.85% and 0.23%, respectively). Using the same peak definition, we applied the analysis to each of the other data sets to evaluate replicability of the coexpression pattern at the xCell, xSubject-xCell and xSubject levels.

PEER analysis: PEER was used by Ling et al. (2024) to identify latent factors (LFs) in the data, akin to using principal components analysis (PCA). Guided by code provided by the authors, we followed their procedure on pseudobulked data (xSubject in our terminology), normalizing and filtering the data as described in their paper and using PEER with default settings and no covariates included in the model, with 10 latent factors to be reported. To be consistent with the rest of our analyses we limited the analysis to protein-coding genes. We confirmed that this had only minor effects (the resulting latent factor has a Speaman’s correlation of 0.849 with that of Ling et al.) (for further details and analyses see the Supplement).

We ran PEER on both the Ling data and the seven studies discussed above using the same preprocessing and filtering steps. To compare PEER LFs across analyses and across data sets, following the procedure of Ling et al., we did not restrict comparisons to a particular ordinal position of the LF and also allowed reversals in the sign. That is, we reported the “best match” LF by absolute Spearman correlation. For example, the original SNAP pattern was latent factor 4 (LF4) as reported by Ling et al. They also performed PEER on their control samples and case samples separately, and reported that this replicated the factor, though was LF1 in controls (Ling et al.’s Supplementary Figure 6B).

## Results

As detailed in Methods, we used seven single-cell studies of human brain for our primary investigations. We also had matched bulk and scRNA-seq samples for the same individuals for two studies. We first describe replicability of three coexpression modes across studies; then preservation across modes for a given study. We then address the biological interpretability of the coexpression patterns. In the last results section, we present an application of our ideas to the analysis of a recently identified coexpression pattern reported to be relevant to schizophrenia (Ling et al., 2024). In the supplement, we further document phenomena that contribute to coexpression in xBulk and its comparability to the other modes.

### Per-cell-type xCell coexpression signals are more replicable across independent single-cell datasets than xSubject

Based on the idea that gene regulation is mostly the same in all humans, we hypothesized that coexpression for a given cell type from a given cohort would be replicable in other cohorts. In contrast, xSubject gene covariance might depend more on the particular subjects. For both of these modalities, we computed odds ratios to quantify the extent that the top 1% edges in each dataset could be reproduced by other datasets (See Methods). In total, we performed 216 comparisons, with one comparison for every pairwise combination within the same cell type. All comparisons yielded odds ratios above 1.0 (mean OR = 37.8; min = 1.01; max = 290.7). For example, 57% of the top 1% coexpression edges in Velmeshev:Inhibitory were recovered by Pineda:Inhibitory (OR = 290.7; q-value < 1e-16, Fisher’s exact test). On average, the level of replicability was highest for inhibitory neurons (mean OR = 87.2) and lowest for OPCs (mean OR = 4.95) (Figure 2A). Although replicability in other cell types was lower, all except one comparison (ROSMAP:OPC vs. Nagy:OPC) were significant at a false discovery rate (FDR) of 0.01. Across cell types, close to 2 million edges were independently discovered in two or more datasets and close to 16,000 were discovered in all seven datasets (Figure 2B).

**Figure 2:**
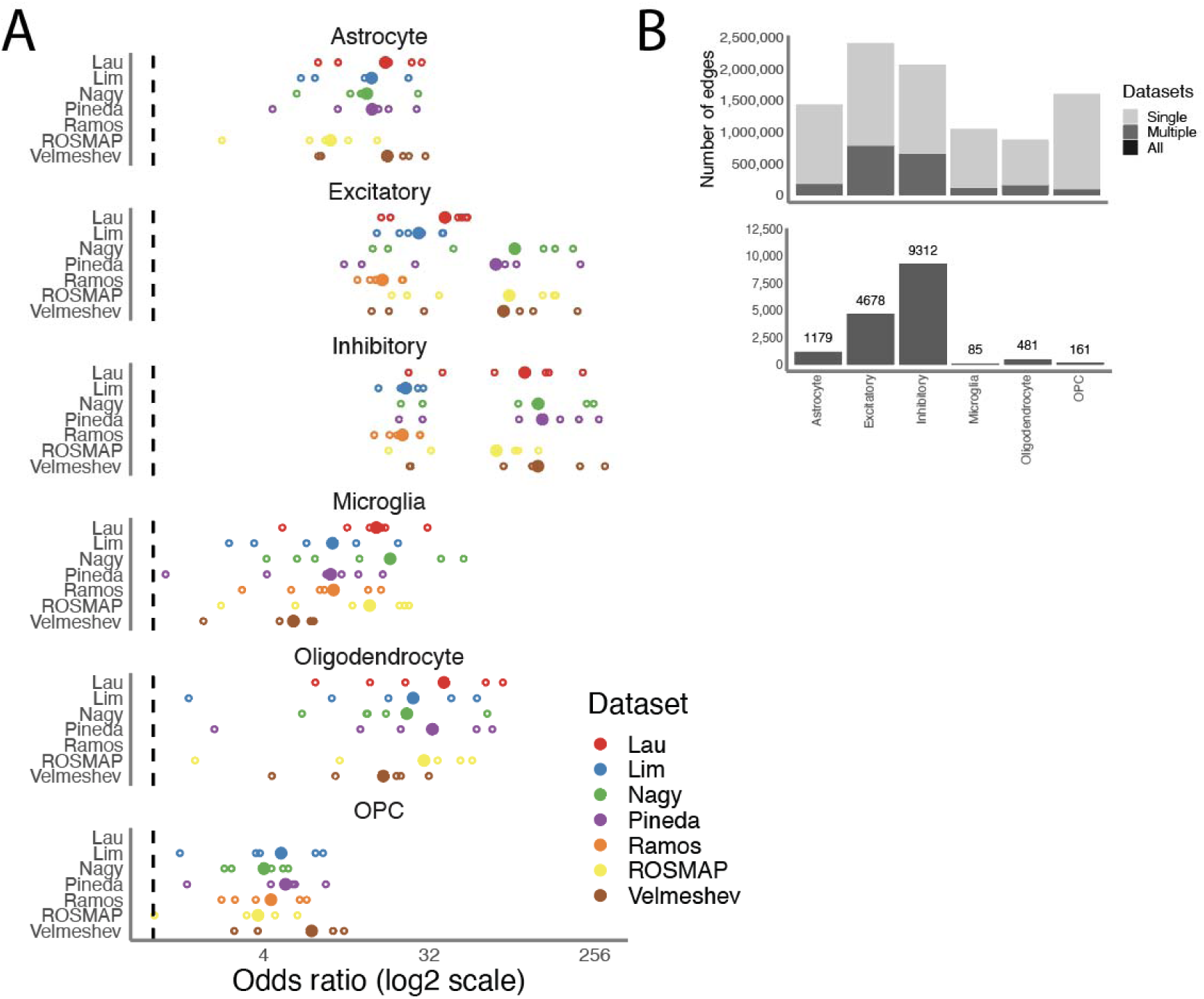
Replicability of xCell coexpression across data sets. A: Each data point shows the odds ratio (x-axis) of a comparison between a reference (y-axis, color) and one test computed using an independent dataset (open data points). The closed, larger data points indicate the average replicability of the given reference compared against all other tests. Odds ratio of 1 is indicated by the vertical dash lines; note the logarithmic scaling. B: Each bar shows th breakdown of coexpression edges that were discovered in all vs. multiple (>2) vs. single dataset for each cell type (x-axis). Data for the edges discovered in all datasets are plotted separately in the bottom panel for legibility.

Repeating this analysis for the xSubject analysis showed that xSubject coexpression patterns are less replicable across studies than xCell (Figure 3A, 3B and 3E). There was an average change in odds ratio of -29.7 (min = - 123.2; max = +1.3) between the xCell and xSubject levels (p-value < 2.55e-10, Wilcoxon signed rank test). We observed the largest difference in replicability in the comparison between Velmeshev:Inhibitory and Pineda:Inhibitory where the odds ratio decreased from 291 at the xCell level to 16.2 at the xSubject level. This corresponded to a drop in edge recovery rate from 57% to 12%. This was consistent in all cell types except OPCs, likely due to the fact that OPCs also had relatively low replicability at the xCell level (Figure 3). Consistent with these observations, few xSubject edges were independently discovered by multiple datasets. For example, in inhibitory neurons, only 10% of all xSubject edges were present in more than one dataset (Figure 3D), compared to 32% at the xCell level (Figure 2B). In the glial cell types, the proportion of singleton dataset discoveries was even higher. Taken together, the replicability of xSubject, though significant, was not nearly as pronounced as that of xCell. Since for a given study xCell and xSubject are constructed from the same data, these findings suggest that differences between studies (i.e. the individuals they studied) may have a disproportionate impact on xSubject compared to xCell coexpression patterns.

**Figure 3.**
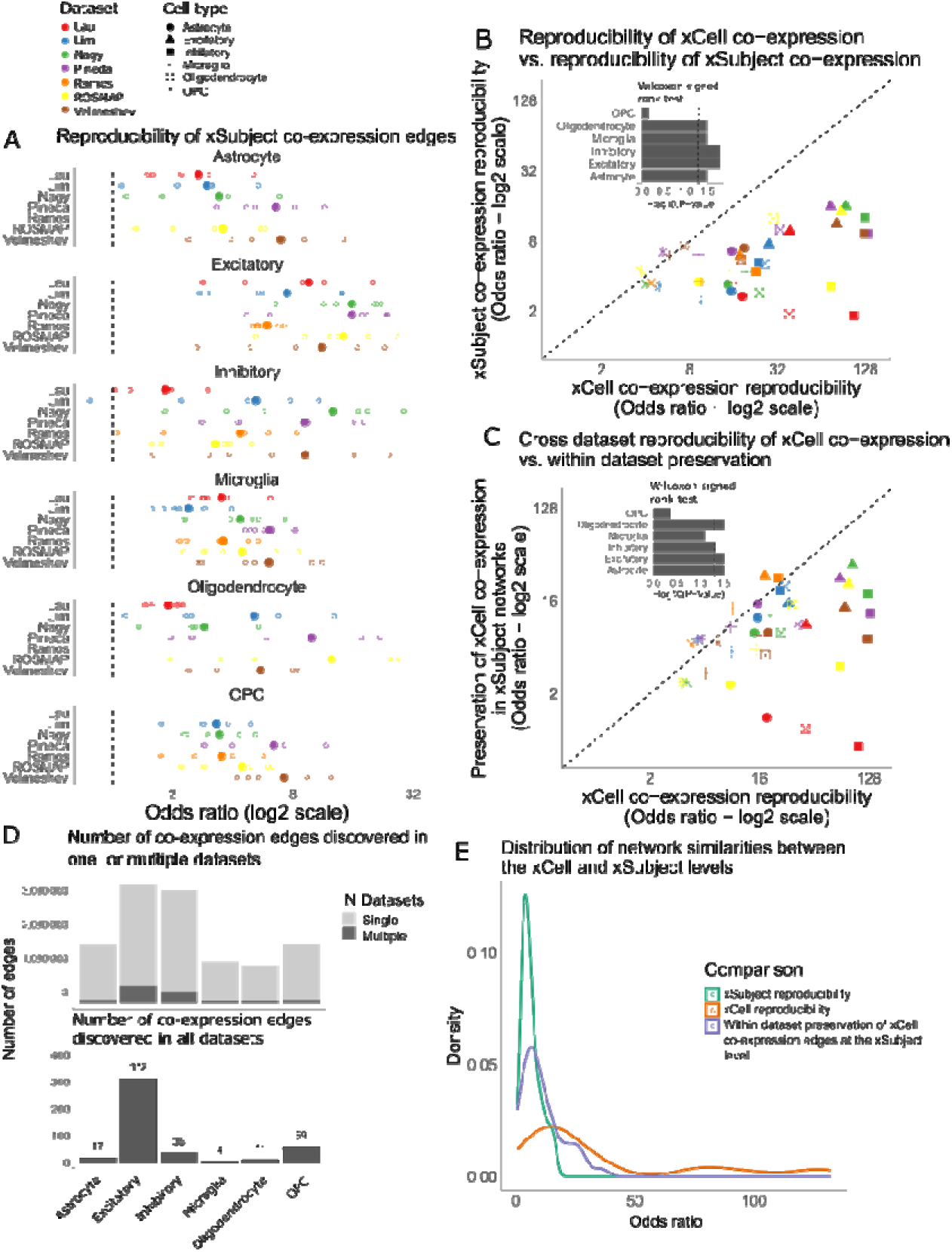
Replicability of xSubject coexpression and preservation of xCell coexpression. (A) Replicability of xSubject coexpression. Each data point shows the odds ratio (x-axis) of a comparison between a reference (y-axis, color) and an independent dataset (open data points). The closed, larger data points indicate the average replicability of the given reference compared against all other test data. Odds ratio of 1.0 is indicated by the vertical dash lines. B) Scatterplot showing replicability at the xCell level (x-axis, summarizing the data from Figure 2A) vs. replicability at the xSubject level (from part A). Each datapoint shows the average odds ratio of comparisons of the indicated reference. The xCell level replicability is significantly higher than the xSubject level replicability across all cell types (main text). Small panel placed within the scatterplot shows the p-values for cell type specific comparisons. Vertical line indicates p-value of 0.05. (C) Scatterplot showing replicability at the xCell level (x-axis) vs. preservation of xCell coexpression edges at the xSubject level for the same reference dataset. The xCell level replicability across datasets is significantly higher than preservation within the same dataset (main text). Small panel placed within the scatterplot shows the p-values for cell type specific comparisons. (D) Each bar shows the breakdown of coexpression edges that were discovered in all vs. multiple (>2) vs. single dataset for each cell type (x-axis). Data for the edges discovered in all datasets were plotted in a separate, bottom panel because they are not visible in the scale of the top panel. (E) Distribution of odds ratios across 39 data groups for the three types of comparisons listed in the legend.

We then compared the coexpression of the two bulk data sets. Velmeshev xBulk was enriched for the coexpression edges in ROSMAP at OR = 47.5. This corresponded to a recovery of 28% of the top 1% edges. The reverse comparison yielded an OR of 36.8 or a recovery of 20% of the top edges in Velmeshev. Though replicability between the two datasets was higher at the xBulk level than at the xSubject level (Figure 4), it was lower compared to xCell replicability for excitatory and inhibitory neurons.

**Figure 4.**
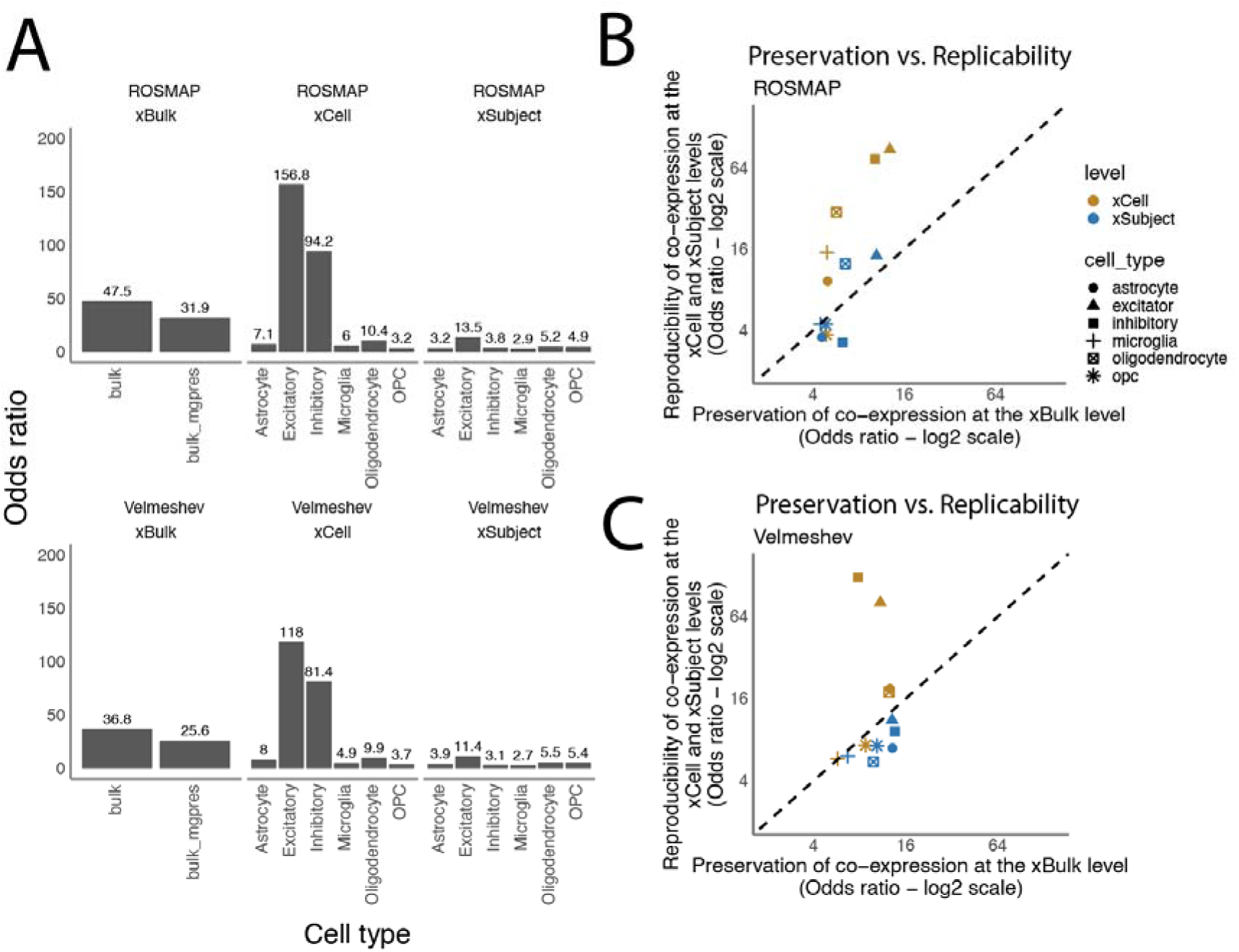
Preservation and replicability of coexpression edges at the xBulk level. (A) Replicability measured between the two datasets at all three levels of analysis, using ROSMAP (top) and Velmeshev (bottom) as the reference dataset. (B) Scatterplot showing the preservation of xCell or xSubject coexpression signals at the xBulk level plotted against xCell or xSubject level replicability in ROSMAP. (C) Same as (B) but for the Velmeshev dataset.

### Many xCell coexpression patterns are not preserved by other analysis modes

Replicability across independent data sets is only one consideration for which coexpression modality to use. Another is the type of biological signal each modality captures. Again, we argue that the analysis most closely aligned to the goal of understanding dynamic regulation is at the xCell level. But if the other modalities contain similar information, there might be little reason to choose one over the other. Based on our modeling studies, we hypothesized that much coexpression observed at the xCell level might be lost if one looks only at the xSubject level due to the absence of explicit cell-to-cell covariation information, and further obscured at the bulk level, which adds in the issues of mixtures of cell types and dilution effects. To test this experimentally, we compared xCell, xSubject and xBulk for each data set (we could only do xBulk for the two data sets for which we had matched scRNA-seq samples). To differentiate this from between-studies replicability, we refer to this as “preservation”: the degree to which coexpression captured by a data set using one mode is also identified by analyzing the same data using the other modes.

We first compared each xCell analysis to the corresponding xSubject coexpression computed from the same data. We detected statistically significant preservation of edges for 37 of 39 comparisons (Figure 3B), with mean odds ratio 11.1 (min = 0.61; max = 35.7), which translated to an average recovery rate of 8.6%. However, these levels of *within dataset* xCell to xSubject preservations were consistently lower than *cross dataset* replicability at the xCell level (Figure 3C). For example, Lau:Inhibitory xCell had an average replicability in other datasets of OR = 106 (recovery rate = 33.5%) but its own preservation at the xSubject level had an OR of 0.61 (recovery rate < 1%). These findings agree with our modeling study and support the hypothesis that xCell coexpression cannot be assumed to be found at the xSubject level and xSubject level coexpression is not simply a proxy for xCell coexpression signals.

We next examined preservation of xSubject coexpression signals at the xBulk level (Figure 4B and C, blue points). This yields just one set of coexpression results (instead of one per cell type as in the other modes), so we compared this to each of the cell-type specific results from xCell and xSubject. In the Velmeshev dataset, the average OR was 11.1 across all six cell types. We observed the highest level of preservation in inhibitory neurons at an odds ratio of 13.6, which corresponded to a recovery of 12% of all xSubject edges. In contrast, microglia had the least preservation at an OR of 6.7 or a recovery of 6% of the xSubject edges. The highest level of preservation in the ROSMAP dataset was in excitatory neurons, at an odds ratio of 10.5, which corresponded to a recovery rate of 9% of the xSubject edges. Microglia had the lowest level of preservation at OR of 4.5 or a recovery of 4% of the xSubject edges. Though the levels of preservation were significantly higher than the null expectation in all cell types (FDR < 0.01), the majority of the xSubject edges were not preserved at the xBulk level.

Next, we evaluated the preservation of xCell patterns in xBulk (Supplementary figure 1D). Notably, in the ROSMAP dataset, xBulk retained substantially more xCell than xSubject signals for both excitatory and inhibitory neurons. Specifically, ROSMAP:xBulk was concordant with edges from excitatory and inhibitory xCell at ORs of 12.8 (recovery rate = 11.1%) and 10.3 (recovery rate = 9.2%) respectively. This was in contrast to enrichment for xSubject edges at ORs of 10.5 (recovery rate = 9.3%) and 6.28 (5.9%). In Velmeshev, oligodendrocytes also had a higher preservation of xCell (OR = 12.5; recovery rate = 11.2%) than xSubject signals (OR = 9.85; recovery rate = 9.0%).

Because bulk tissue contains no direct information about cell-to-cell variability, it may seem surprising that there is higher concordance of xCell coexpression with it than xSubject coexpression, especially since bulk tissue contains multiple cell types. We hypothesized that might in part be explained by the effects of cell type composition variance (CCV), on the basis of our simulation study and previous work. CCV can be a major source of coexpression signal at the xBulk level (Farahbod and Pavlidis (2020)), inducing apparent coexpression among genes that have similar cell type expression profiles. We predicted that correcting for CCV effects should reduce concordance with xCell without compromising concordance with xSubject. We present extensive investigation of this hypothesis in the Supplement. To summarize, we found that corrections for CCV indeed generally reduces preservation of xCell patterns in xBulk (Extended Data Figure 1). This suggests that sometimes differences *between* cell types can agree with differences *among* cells of a given type. There are also interesting exceptions that survive CCV correction, which can be attributed to cases where dilution effects are avoided by the pattern being present in all cell types. Furthermore, we identified strong evidence for the effects of dilution in bulk tissue on the recovery of patterns apparent at the xSubject or xCell level (Extended Data Figure 4).

### Curated regulatory interactions are enriched in cell-level coexpression patterns

Comparison of coexpression at the different levels of analysis informs their concordance and differences. In theory, xCell coexpression would be closest to coexpression within single cells, which is the actual target of the analysis. As such, we hypothesized xCell would be most enriched for signals of dynamic regulation. This is difficult to assess, as there are limited “gold standards”, all with heavy biases and do not define negatives (a lack of regulation of a given gene by a given TR), nor necessarily distinguish between “static” and “dynamic”. An additional problem is most databases of TR-target relations lack information on the cellular context. To limit the latter problem, we used a catalogue of literature-curated regulatory interactions which has such annotations (Chu et al., 2021) though the other biases are still present and should be kept in mind as a limitation. We defined c - regulated genes to be pairs of genes that share a common upstream transcription factor according to Chu et al. (2021) and which were annotated to be relevant to the brain context, yielding 1,480 pairs. We measured the performance of xCell, xSubject, and xBulk coexpression in prioritizing pairs of co-regulated genes, summarized as AUROC (See Methods).

We found significant enrichment for co-regulated pairs in the xCell coexpression analyses. Seven of the 39 were significant at p < 0.05 (Mann-Whitney U test) and four remained significant after multiple test correction (FDR < 0.1). Although most of the xCell analyses did not meet the threshold for statistical significance, the majority (30 of 39) had AUROCs > 0.5, suggesting some level of enrichment above random chance (Figure 5). In particular, all of the excitatory neuron xCell analyses had AUROCs > 0.5. We further compared these results to shuffled TR-target pairs, representing an empirical null (see Methods). Ten of the 39 xCell data sets were significant at p-value < 0.05 using this method, indicating that the enrichment performance was specifically driven by the co-regulatory gene pairs (Figure 5A).

**Figure 5.**
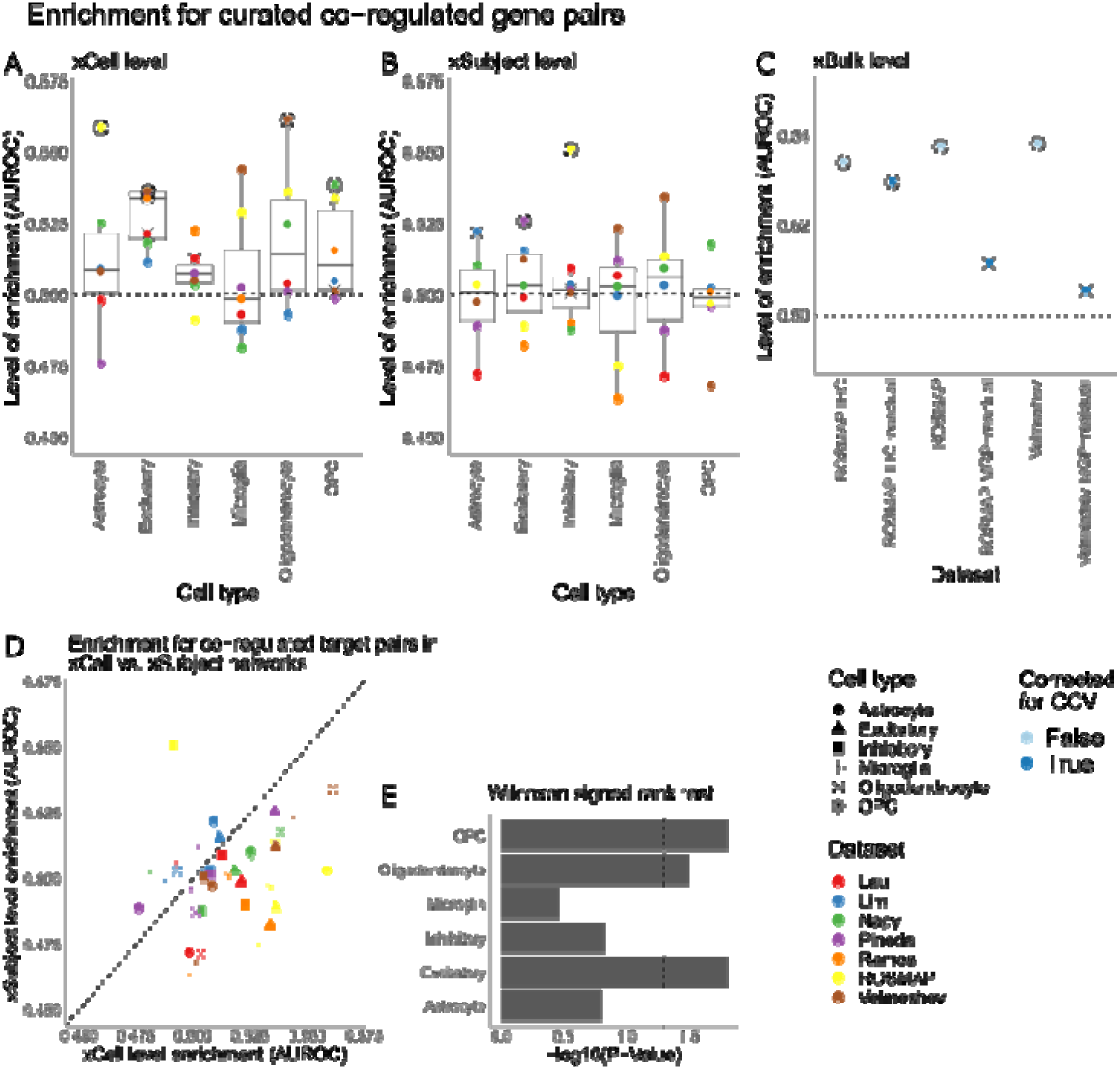
Recovery of curated regulatory interactions. (A) Each data point shows the level of enrichment (AUROC, y-axis) of xCell coexpression for a given cell type (x-axis) and dataset (color). Significant values at p-value < 0.05, computed using the Mann-Whitney U test are circled. Significant values at empirically derived p-value < 0.5 (See main text) is crossed. (B) Same as (A) but for xSubject level coexpression. (C) Same as (A) but for xBulk level coexpression. Importantly, the enrichment in the CCV corrected data for all three corrective procedures are also plotted. (D) Each datapoint is a data group. The level of enrichment for co-regulated genes at the xCell level (x-axis) for the given data group are plotted against the level of enrichment in the corresponding xSubject analysis (y-axis). The xCell level enrichment is significantly higher than the xSubject level replicability across all cell types (main text). (E) Bar plots showing the p-values for the difference between xCell vs. xSubject enrichment within each cell type shown in (D).

Although the levels of enrichment measured using AUROC were only slightly higher than 0.5, they were on par with previous assessments of coexpression (Garcia-Alonso et al., 2019). The low performance on an absolute scale was also expected due to the limited number of curated targets. However, the quality of the coexpression likely plays a role as well. Consistent with our observation of lower replicability, microglia also had the lowest AUROCs. Despite these limitations, there was a level of detectable enrichment for the literature curated regulatory interactions.

A key question is how xSubject and xBulk coexpression performs relative to xCell. We found that the xCell coexpression performed better than xSubject (Figure 5B). Only two of the 39 were nominally significant in xSubject (p < 0.05) and five were significant when compared to the empirically derived null distributions. Only 23 of the 39 had AUROCs > 0.5 (mean = 0.50). Overall, AUROC decreased by up to 0.06 between xCell and xSubject levels (p-value < 1.7e-4, Wilcoxon signed rank test; Figure 5E). The reduction in enrichment was significant for three out of six cell types.

Finally, at the xBulk level, both ROSMAP and Velmeshev datasets had significant levels of enrichment for co-regulated gene pairs (Figure 5C), which exceeded that in xSubject. We hypothesized that this elevated performance may be partly due to CCV. Accordingly, enrichment in both datasets decreased with CCV correction (Figure 5C). This indicates that static cell type differences may have meaningful contributions to the enrichment of co-regulated gene pairs, but we believe this is conceptually, and likely phenomenologically, different than coexpression observed at the xCell level.

Overall this assessment suggests that enrichment for known regulatory interactions can be detected at all levels of coexpression analysis. This is consistent with the literature, as researchers have been using these modes for regulatory inference and reporting some success. The fact that xCell performs at least comparably to the other modes on this limited benchmark is encouraging. The other advantages of xCell, including high replicability across studies and more direct connection to sought-after intracellular signals of regulation are at least as important in our view.

### Case study: The “SNAP” pattern

We’ve laid out a case that different coexpression analysis modes will yield different results in ways that can affect interpretation and replicability. In this section we delve into a particular example. (Ling et al., 2024) analyzed 191 post-mortem prefrontal cortex samples from controls and people with schizophrenia with 10X Chromium snRNA-seq. They conducted two analyses that relate to our questions. First, they used a version of the xSubject approach to identify a pattern they called “SNAP” (“synaptic neuron and astrocyte pattern”). To attain this they used PEER (Stegle et al., 2010), a method that (as applied by Ling et al.) in effect identifies clusters of coexpressed genes (see Methods and Ling et al. 2024). SNAP was identified as a PEER latent factor (LF4) that had a difference in factor loadings in schizophrenia and also with age. The top 1000 genes with the highest contributions to the pattern were referred to as the “strongest” SNAP genes and described as “strong[ly]” coexpressed (Ling et al., 2024). Second, they used a version of the xCell-xSubject approach. They did this separately on two cell types (excitatory neurons and astrocytes) in contrast to the more common practice which uses all cells. Ling et al. did this out of concern for an issue we have highlighted: that cell type differences might dominate the coexpression patterns, over patterns present within a cell type. They reported some concordance (preservation, in our terms) between the xSubject and xCell-xSubject patterns with respect to SNAP. However, as far as we know, Ling et al. did not attempt to isolate the effects of subject-to-subject variation from cell-to-cell, which is one of our main concerns. Ling et al. also did not assess whether the SNAP pattern is present in other data sets (replication). It is reasonable to hypothesize it is present in other data sets as Ling et al. interpreted the pattern as potentially reflecting a fundamental feature of the human brain, not a phenomenon specific to their data set, and it was present in controls as well as schizophrenia cases. As a preliminary step, we confirmed that we could reproduce their key PEER result (see Methods and Supplementary Figure 3). We then conducted analyses of the reported top genes in the SNAP pattern (1000 genes selected by magnitude of the loading as per Ling et al.), as well as additional PEER analyses.

Using our xSubject approach on the Ling data, we observed coexpression of many of the SNAP genes in excitatory neurons and astrocytes, generally validating their findings on their own data with a different approach. This was much clearer for the genes which had positive loadings in the SNAP factor (Supplementary Figure 4, right-hand panels). We note that few of the SNAP pairs meet the top 1% coexpression criterion that we used in the previous analysis (nor did Ling et al. claim they would be), but many are still above the background expectation. We also noted that the ribosomal protein coding genes show much less xSubject coexpression in Ling’s excitatory neurons, compared to what we observed in other data groups, and for Ling’s astrocytes (Supplementary Figure 4).

We then conducted coexpression using two coexpression modes that use cells as the unit of analysis and examined preservation of SNAP across these levels. First, we used xCell on astrocytes and separately on excitatory neurons, using our aggregation approach. Then, we mimicked Ling et al.’s second analysis by conducting xCell-xSubject coexpression on the excitatory neurons and separately on astrocytes. Coexpression of the SNAP genes was overall lower at the xCell level and the distribution of correlations shows that the genes tend to “split” into two clusters that are negatively correlated with each other (Supplementary Figure 4). The xCell-xSubject pattern showed much stronger agreement with xSubject (Figure 6A), consistent with Ling’s interpretation that subject-to-subject variability drives the pattern. This difference of modes is supported by the lack of strong ribosomal protein gene coexpression in Ling’s excitatory neurons at the xSubject level, in contrast to the xCell level (Supplementary Figure 4), a distinct pattern from what we observed with the other data sets we examined. Overall, this shows that while some of the SNAP pattern is preserved across modalities (even xCell) in the Ling data, many features of the coexpression patterns are affected.

**Figure 6:**
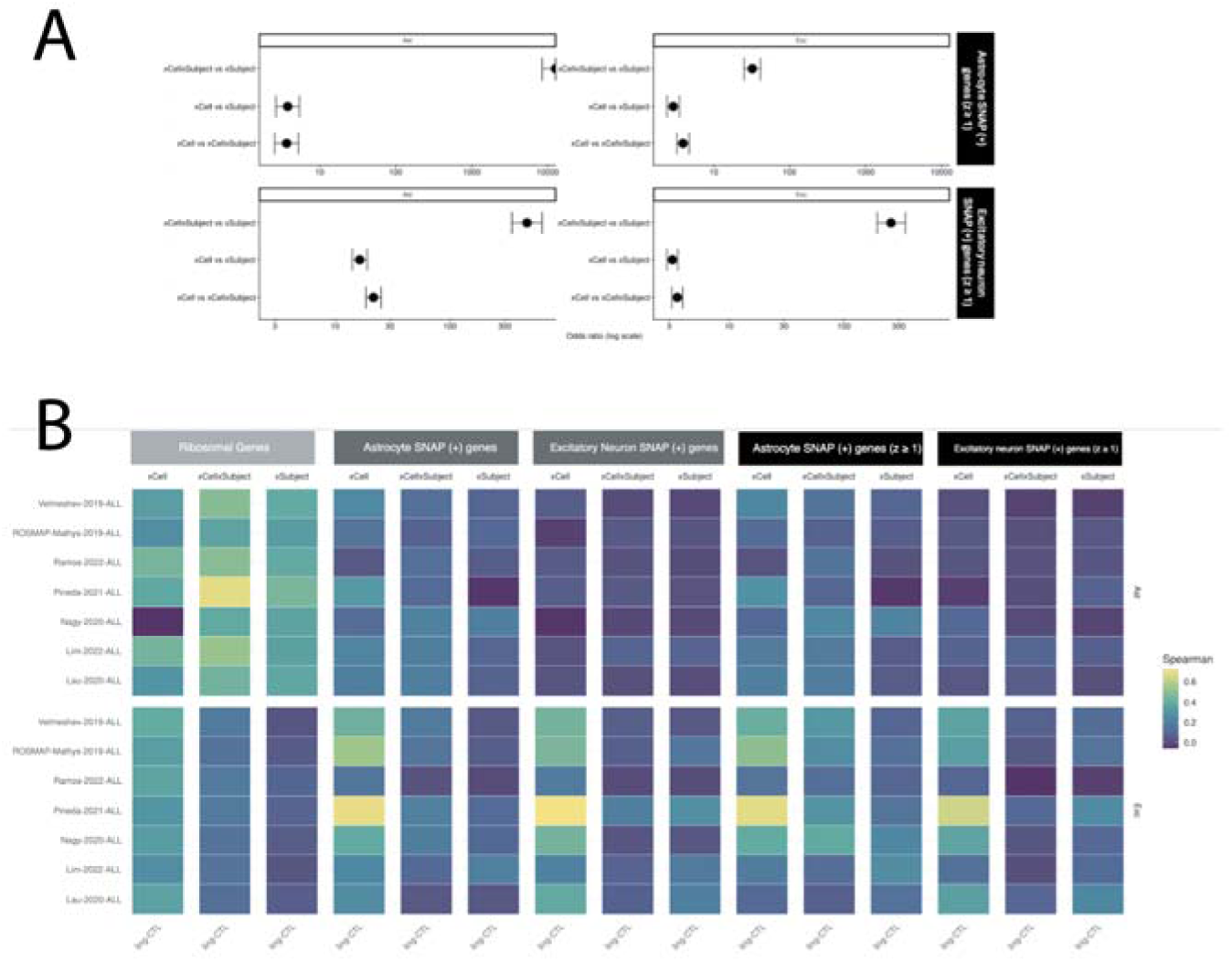
Analysis of the SNAP pattern from Ling et al. (2024). A: Odds ratios for preservation of SNAP coexpression at different levels of analysis. B: Patterns of replication of SNAP coexpression in the other seven data sets considered. Replicability was higher when correlations were computed at the xCell level than at xSubject. In astrocytes, xCell correlations showed higher replication (mean Spearman ρ = 0.17 ± 0.09, range 0.02–0.28) than xSubject correlations (mean ρ = 0.06 ± 0.08, range −0.05–0.22). In excitatory neurons, xCell replication was also higher (mean Spearman ρ = 0.33 ± 0.18, range 0.09–0.64) compared with xSubject-level replication (mean ρ = 0.14 ± 0.10, range −0.03–0.26). Pairwise comparisons between analysis modes were evaluated using the Wilcoxon ranksum test with Bonferroni correction (xCell vs xCell×Subject: adjusted p = 0.001; xCell vs xSubject: adjusted p = 1.4 × 10⁻□; xCell×Subject vs xSubject: adjusted p = 0.389).

We next tested whether patterns found xSubject, xCell or xCell-xSubject in the Ling data are replicated in the other brain data sets. Overall, the coexpression of the SNAP genes was not well reproduced at the xSubject and xCell-xSubject levels in other studies (Figure 6B). As a further check of replicability, we applied the PEER methodology to the other data sets, and sought the latent factor that most resembles SNAP in terms of gene loading correlations (see Methods). The SNAP pattern was not clearly identified in any of them; the median best match had a Spearman’s correlation of 0.15 with Ling’s SNAP loadings, with the highest correlation being 0.33 with LF3 in the Lim data set (out of a total of 210 comparisons; Supplemental Figure 3C). The 1000 top-loaded genes in Lim’s LF3 only contains 84 of the top genes identified by Ling et al., and of those only 17 had the same cell type association as they had in Ling et al.

We hypothesized that the subset of xSubject SNAP coexpression that we considered preserved in the Ling data at the xCell level (1,929 edges in astrocytes and 1,482 edges in excitatory neurons) would be more replicable in xCell coexpression of other data sets. Indeed this is the case, replication of SNAP coexpression was stronger when correlations were computed at the xCell level than when computed at subject-resolved levels (Figure 6B). This trend was even stronger for the subset of SNAP edges that also met the top 1% coexpression criterion. (Figure 6B).

## Discussion

We found that there is limited direct preservation of coexpression signals across analyses at different levels of resolution. The xCell coexpression patterns were largely distinct from xSubject coexpression. The mixture of different cell types at the bulk level results in even more distortion of coexpression signals due to dilution effects as well as additional coexpression patterns induced by composition variation (CCV). There is a subset of genes with exceptional levels of preservation of coexpression in bulk data, and we have documented their distinguishing characteristics, which include coincidence of CCV-induced coexpression or the presence of inter-cell type synchrony.

The benefit of xCell analysis is supported by its high replicability across studies, enrichment for literature curated co-regulated gene pairs, as well as its interpretability. Our results also suggest that sequencing depth, number of cells and subjects is probably important to a successful and replicable coexpression analysis. For example, microglia and OPCs were relatively data-sparse and also less replicable. This suggest that, with current technology, large amounts of data can be required to obtain robust and replicable xCell coexpression patterns, a conclusion further supported by our report in which we used (non-cell-type-specific) xCell patterns to closely examine coexpression patterns of transcription factors (Morin et al., 2025).

We found that xSubject coexpression was less replicable across studies than xCell and was notably different from xCell coexpression patterns extracted from the same data. We emphasize that the xCell coexpression patterns extracted from the exact same data sets were much more replicable, serving as a positive control that there isn’t just a data quality issue. We did identify a small number of clusters, including the ribosomal protein genes, that have coexpression patterns that tend to be observed no matter the analysis mode. But they are the exception. Our current interpretation is that some xSubject coexpression is specific to the particular set of subjects in a study, that is, it is idiosyncratic.

When it comes to inferring regulation such as targets of a TF, because xSubject covariation is driven by differences among subjects, there has to be sufficient variation in the activity of the TFs across individuals for their targets to be detectably coexpressed. If the activity of a TF is too stable across subjects, its targets would not show enough variation for coexpression to be detected. Our results suggest that this might dominate. We note that pseudobulking per subject (per cell type) appears to be less commonly used in single-cell genomics studies of coexpression and GRN inference methods but is routinely used in “bulk” analyses of purified cells or cell lines. It is important to stress that xSubject can still potentially capture intracellular regulatory signals. Studies that use synchronized cell cycle states in cultured cells (only one cell type) analyzed in bulk are a successful application of this principle (Eisen et al., 1998; Spellman et al., 1998).

Our case study of the SNAP coexpression pattern supports our conclusions. Within the Ling et al. data set, preservation of the patterns from the xSubject level (used to discover SNAP) to the xCell level was limited, consistent with our observations of other data sets. The xSubject SNAP pattern was also not reproduced in other data sets. The reasons for this are not certain but an interpretation consistent with our other findings is that xSubject observations like SNAP are vulnerable to not being generalizable to different sets of subjects.

Furthermore, consistent with what we observed for other data, cell-level coexpression patterns in the Ling data (xCell, not driven by subject-to-subject variance) have a better chance of being replicable across studies. While a subset of the xSubject SNAP pattern shows preservation at the xCell level, if Ling et al had started with xCell, it would likely have led to a different and potentially more replicable pattern being found. Future searches for disease-associated coexpression patterns should consider these issues carefully.

We were interested to find significant preservation of xCell signals in xBulk coexpression. A deeper analysis revealed that much of the xBulk coexpression (and its agreement with xCell) was due to CCV. Because CCV is a proxy for stable cell type differences, this suggests that co-regulated genes (within a cell type) can share cell-type specificity, which isn’t too surprising and agrees with the interpretation of (Harris et al., 2021). But clearly xCell and xBulk are not interchangeable, and one should be especially wary of interpreting xBulk coexpression directly as signals of dynamic regulation. If one is interested in static cell type differences, differential expression analysis between cell types, when feasible, might be preferred over trying to extract this information indirectly from heterogeneous bulk samples. On the other hand, there were some xBulk coexpression signals that appear to represent cases of signal propagation from the xSubject level. Our analysis shows inter-cell type expression synchrony is usually required for this to happen. That is, the coexpression pattern needs to be present in all cell types, so that they can still emerge when the cell types are mixed. But without such prior knowledge of synchrony, distinguishing such patterns in bulk tissue from those due to CCV or other sources of xSubject variability is probably not possible.

In summary, in agreement with our simulation study, we have shown that there is limited preservation of xCell coexpression across the data resolutions, despite its high replicability across studies, with notable exceptions for which we are able to provide some explanation. While our analyses are based on studies of human brain, we believe the general principles we have outlined are likely to hold for other situations. We therefore recommend that efforts to look at cell-type-specific dynamic intracellular regulation are best done using xCell analysis from multiple biological replicates. The aggregation approach we apply here is a natural and apparently effective way to do this, but other approaches can be envisioned (Farahbod and Pavlidis, 2020). Coexpression studies that use xBulk data should be carefully interpreted in light of the role of cellular composition variation and the impact of dilution effects, and attending to the distinction between xCell and xSubject effects. Overall we hope our study contributes a stronger conceptual basis for coexpression analysis decisions.

## Author contributions

CPC: Designed and performed analyses, drafted the manuscript and figures; NE, RX: additional data analysis, contributed to writing and figures; JG, BX: contributed to data assembly and analysis; AM: contributed to project conception and development of the approach; PP: Project conception, supervision and writing.

## Acknowledgements

We thank the authors who made data available for our analyses. The results published here are in part based on data obtained from the AD Knowledge Portal (https://adknowledgeportal.org). Samples for that data were provided by the Rush Alzheimer’s Disease Center, Rush University Medical Center, Chicago. Data collection was supported through funding by NIA grants P30AG10161, R01AG15819, R01AG17917, R01AG30146, R01AG36836, U01AG32984, U01AG46152, the Illinois Department of Public Health, and the Translational Genomics Research Institute. We thank Drs. Emi Ling and Steven McCarroll for assistance with their data. We acknowledge Guillaume Poirer-Morency, Dmitry Vavilov and Sanja Rogic for technical and administrative support.

## References

Aibar S, González-Blas CB, Moerman T, Huynh-Thu VA, Imrichova H, Hulselmans G, Rambow F, Marine J-C, Geurts P, Aerts J, van den Oord J, Atak ZK, Wouters J, Aerts S (2017) SCENIC: single-cell regulatory network inference and clustering. Nature Methods 14:1083–1086 Available at: https://www.nature.com/articles/nmeth.4463 [Accessed February 15, 2018].

Chu CP, Morin A, Pavlidis P (2025) Limits to the inference of gene regulation from bulk tissue expression data. :2024.10.24.619521 Available at: https://www.biorxiv.org/content/10.1101/2024.10.24.619521v3 [Accessed October 2, 2025].

Chu EC-P, Morin A, Chang THC, Nguyen T, Tsai Y-C, Sharma A, Liu CC, Pavlidis P (2021) Experiment level curation of transcriptional regulatory interactions in neurodevelopment. PLOS Computational Biology 17:e1009484 Available at: https://journals.plos.org/ploscompbiol/article?id=10.1371/journal.pcbi.1009484 [Accessed October 20, 2021].

Eisen MB, Spellman PT, Brown PO, Botstein D (1998) Cluster analysis and display of genome-wide expression patterns. Proc Natl Acad Sci U S A 95:14863–14868 Available at: http://www.ncbi.nlm.nih.gov/pmc/articles/PMC24541/ [Accessed August 5, 2014].

Farahbod M, Pavlidis P (2019) Differential coexpression in human tissues and the confounding effect of mean expression levels. Bioinformatics 35:55–61 Available at: https://academic.oup.com/bioinformatics/article/35/1/55/5048456 [Accessed March 19, 2019].

Farahbod M, Pavlidis P (2020) Untangling the effects of cellular composition on coexpression analysis. Genome Res 30:gr.256735.119 Available at: http://genome.cshlp.org/lookup/doi/10.1101/gr.256735.119 [Accessed July 5, 2020].

Garcia-Alonso L, Holland CH, Ibrahim MM, Turei D, Saez-Rodriguez J (2019) Benchmark and integration of resources for the estimation of human transcription factor activities. Genome Res 29:1363–1375 Available at: https://www.ncbi.nlm.nih.gov/pmc/articles/PMC6673718/ [Accessed October 7, 2019].

Harris BD, Crow M, Fischer S, Gillis J (2021) Single-cell co-expression analysis reveals that transcriptional modules are shared across cell types in the brain. Cell Systems Available at: https://www.sciencedirect.com/science/article/pii/S2405471221001538 [Accessed May 25, 2021].

Langfelder P, Horvath S (2008) WGCNA: an R package for weighted correlation network analysis. BMC Bioinformatics 9:559 Available at: http://www.biomedcentral.com/1471-2105/9/559/abstract [Accessed November 12, 2014].

Ling E et al. (2024) A concerted neuron–astrocyte program declines in ageing and schizophrenia. Nature:1–8 Available at: https://www.nature.com/articles/s41586-024-07109-5 [Accessed March 7, 2024].

Marbach D, Prill RJ, Schaffter T, Mattiussi C, Floreano D, Stolovitzky G (2010) Revealing strengths and weaknesses of methods for gene network inference. PNAS 107:6286–6291 Available at: http://www.pnas.org/content/107/14/6286 [Accessed April 9, 2013].

Morin A, Chu CP, Pavlidis P (2025) Identifying reproducible transcription regulator coexpression patterns with single cell transcriptomics. PLoS Comput Biol 21:e1012962 Available at: 10.1371/journal.pcbi.1012962.

Pratapa A, Jalihal AP, Law JN, Bharadwaj A, Murali TM (2020) Benchmarking algorithms for gene regulatory network inference from single-cell transcriptomic data. Nat Methods 17:147–154.

Speir ML, Bhaduri A, Markov NS, Moreno P, Nowakowski TJ, Papatheodorou I, Pollen AA, Raney BJ, Seninge L, Kent WJ, Haeussler M (2021) UCSC Cell Browser: visualize your single-cell data. Bioinformatics 37:4578–4580 Available at: 10.1093/bioinformatics/btab503 [Accessed July 3, 2025].

Spellman PT, Sherlock G, Zhang MQ, Iyer VR, Anders K, Eisen MB, Brown PO, Botstein D, Futcher B (1998) Comprehensive Identification of Cell Cycle–regulated Genes of the Yeast *Saccharomyces cerevisiae* by Microarray Hybridization Fink GR, ed. MBoC 9:3273–3297 Available at: https://www.molbiolcell.org/doi/10.1091/mbc.9.12.3273 [Accessed March 22, 2023].

Stegle O, Parts L, Durbin R, Winn J (2010) A Bayesian Framework to Account for Complex Non-Genetic Factors in Gene Expression Levels Greatly Increases Power in eQTL Studies. PLoS Comput Biol 6:e1000770 Available at: 10.1371/journal.pcbi.1000770 [Accessed February 4, 2011].

