## Supplementary Information for "Evaluation of analysis modes for RNA coexpression in single-cell and bulk tissue"

**Supplementary methods, text, tables, and figures**

### List of supplementary tables and figures

Supplementary Table 1: Single cell datasets used in the primary analyses

Supplementary Figure 1: Properties of the scRNA-seq data sets, supporting their general comparability

Supplementary Figure 2. Schematic of the xCell analysis approach as applied to one cell type in one data set.

Supplementary Figure 3: PEER analysis of SNAP

Supplementary Figure 4: Coexpression of the strongest SNAP-associated genes as reported by Ling et al. (2024), in the Ling data

### List of extended data figures

Extended Data Figure 1. Correction for composition variation reduces preservation of xCell signals at the xBulk levels

Extended Data Figure 2: Relationship between gene variation across cell types and MGP-estimated CCV effects for the Velmeshev data set.

Extended Data Figure 3: Preservation of some clusters is driven by CCV effects:

Extended Data Figure 4 The effects of dilution of cell type specific xSubject expression patterns

Extended Data Figure 5. Inter-cell type synchrony of ribosomal protein genes (RPGs).

Extended Data Figure 6. Association between inter-cell type synchrony and preservation of xSubject co-expression patterns in xBulk (Velmeshev)

Extended Data Figure 7 Association between inter-cell type synchrony and preservation of xSubject co-expression patterns in xBulk networks (ROSMAP)

**Supplementary Table 1. Single cell datasets used in the primary analyses**

| **Study** | **Region** | **N. samples** | **N. cells** | **Library size (mean UMIs/cell)** | **Age** | **Condition** |
| --- | --- | --- | --- | --- | --- | --- |
| (Lau et al., 2020) | Prefrontal cortex | 21 | 169,496 | 4,267 | 60 – 95 | Alzheimer’s disease |
| (Lim et al., 2022) | Cingulate cortex; Nucleus accumbens; caudate nucleus | 32 | 72,115 | 7,111 | Unknown | Huntington disease |
| (Nagy et al., 2020; Pineda et al., 2024) | Dorsolateral prefrontal cortex | 36 | 78,886 | 2,987 | 18 – 87 | Major depressive disorder |
| (Pineda et al., 2024) | Primary motor cortex | 66 | 380,610 | 8,142 | 50 – 90 | Amyotrophic lateral sclerosis; frontotemporal lobar degeneration |
| (Ramos et al., 2022) | Prenatal germinal cortex and cortical plate; adult subventricular zone and posterior frontal cortex | 36 | 157,898 | 6,467 | 17 – 41 gestational weeks; adult | None |
| (Mathys et al., 2019) (ROSMAP) | Prefrontal cortex | 48 | 70,634 | 2,862 | 75 – 90+ | Alzheimer’s disease |
| (Velmeshev et al., 2019) | Prefrontal cortex, anterior cingulate cortex | 41 | 104,559 | 5,275 | 4– 22 | Autism spectrum disorder |

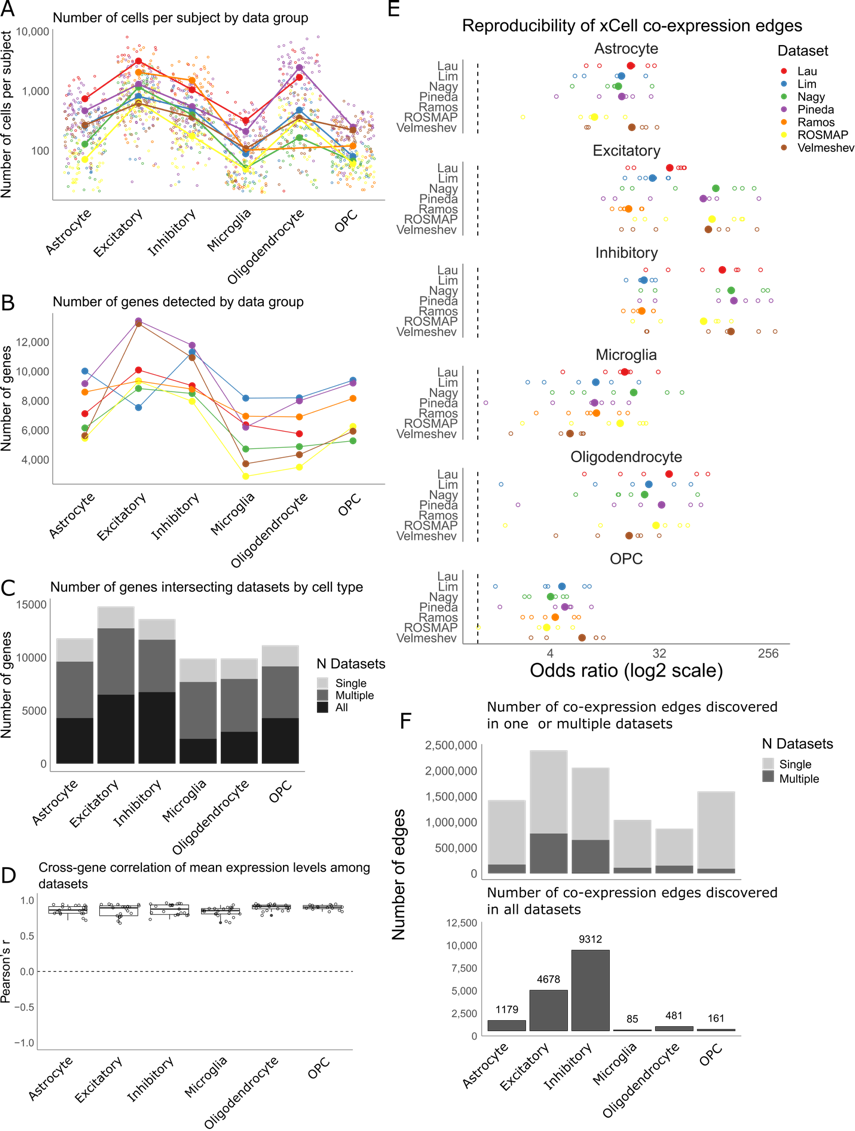

**Supplementary figure 1: Properties of the scRNA-seq data sets, supporting their general comparability** (A) The number of cells per subject are shown on the y-axis, against cell type on the x-axis. Each small datapoint is a subject, colored by dataset. The large datapoints show data group-level averages. Lines connect points of the same dataset across cell types. (B) Genes retained were variable across data groups. Each data point shows the number of genes (y-axis) detected in each data group. The data points are organized by cell type (x-axis) and colored by dataset. (C) The genes detected were highly consistent across different datasets for the same cell type. Each bar shows the breakdown of genes by detection in all vs. multiple (>2) vs. single datasets. (D) Mean expression levels were highly concordant across different datasets for the same cell types. Each data point shows the correlation of expression across the intersecting genes between one pair of datasets.

**Figure** Error! No text of specified style in document.**‑1. Construction of xCell coexpression by aggregation.**

Schematic showing the construction of xCell coexpression by aggregating subject level coexpression matrices. The cell level expression matrix for each subject is first used to compute subject-specific xCell coexpression by computing coexpression across single cells between all pairs of genes. Following rank transformation, a consensus is subsequently derived by averaging all subject level matrices in the given data group (e.g. for a given cell type).

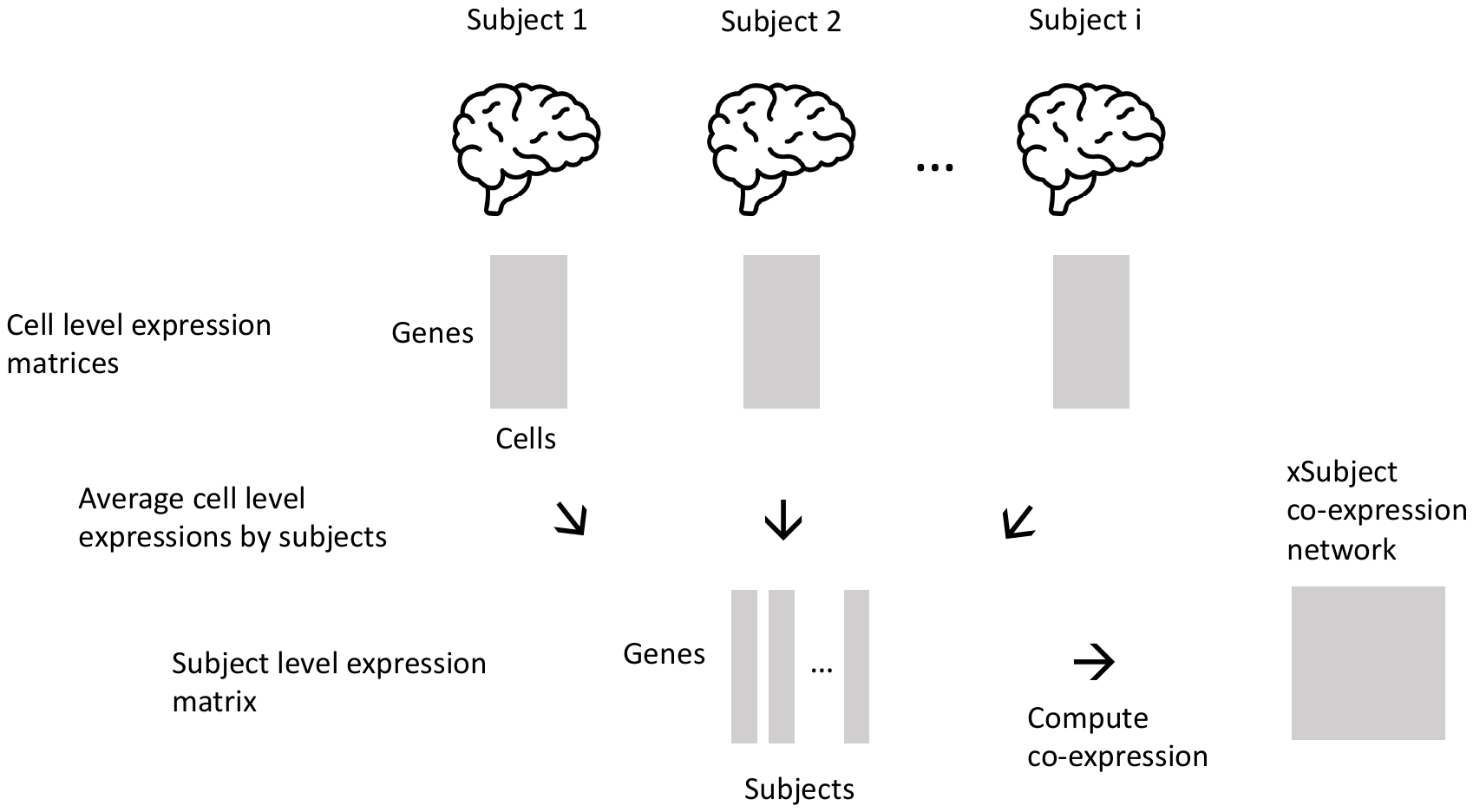

**Figure** Error! No text of specified style in document.**‑2. Construction of xSubject coexpression.**

Schematic showing the construction of a xSubject coexpression by first deriving subject level expression profiles, for a given cell type. First, the cell level expression matrix, where the rows are genes and the columns are cells, for each subject is collapsed into a single column or vector by averaging the cells (pseudobulking). The subject level expression vectors are then concatenated horizontally to form a new expression matrix, which is in turn used to compute the xSubject coexpression. This is done separately for each cell type.

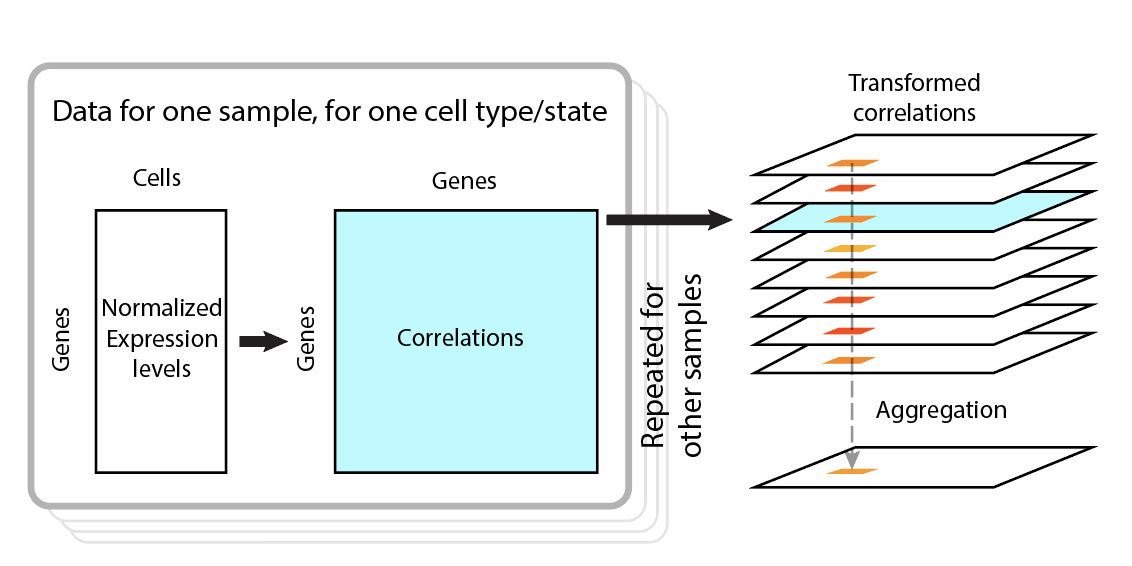

**Supplementary figure 2. Schematic of the xCell analysis approach as applied to one cell type in one data set.** Coexpression matrices are computed for each sample (left side of figure), rank transformed, and aggregated to a final (normalized) ranking for each gene-gene pair (right side). The end result is that the correlation between a given pair of gene is summarized as an average of the rankings among the samples (orange-red squares). This process is conducted independently for each cell type, and for each data set.

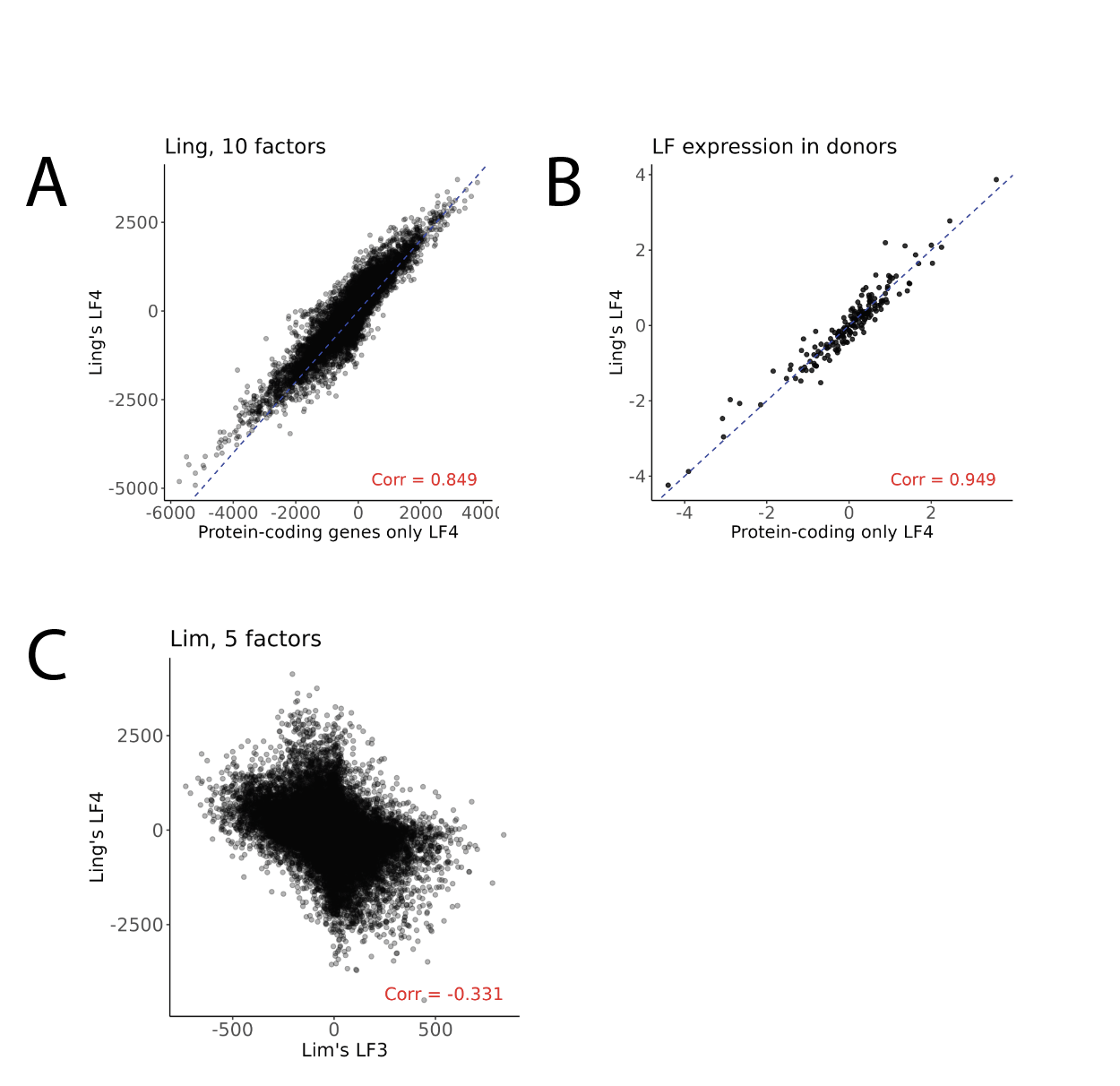

Supplementary Figure 3: PEER analysis. A: Confirmation of recovery of the SNAP pattern (Ling’s Latent Factor 4, LF4) using PEER on the Ling data set (10 factors sought). The main difference is that we limited the data set to protein-coding genes. B: Subject loadings in the Ling data set compared between our analysis and Ling’s. C: The PEER factor among all the other data sets we examine that has the highest correlation with SNAP (Lim LF3, 5 factors sought).

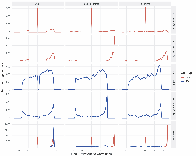

**Supplementary Figure 4: Coexpression of the strongest SNAP-associated genes as reported by Ling et al. (2024), in the Ling data.** Each column represents a level of analysis (xCell, xCell-xSubject and xSubject from left to right). Each of the first four rows of plots is for the SNAP genes that were associated with a particular cell type (astrocytes [ast], or excitatory neurons [exc]), with matching sign of the PEER latent factor 4 loading (positive and negative; numbers of genes indicated by *n*). The last row shows the pattern for ribosomal protein genes, with both cell types plotted. Each plot is a histogram of the Z-score transformed correlations of pairs of genes. If all the SNAP genes are highly coexpressed as claimed by Ling et al. (which should tend to be the case for genes sharing loading sign), we would expect a peak at the right-side of the distribution. The presence of peaks both at positive and negative scores (e.g. for astrocyte-associated SNAP genes in the xCell analysis) indicates a mixture of positive and negative correlations among the pairs, indicating the genes break into at least two anti-correlated patterns. A peak at zero indicates pairs that show no coexpression. In contrast, a more uniform distribution without a pronounced peak at either extreme suggests that the genes are not generally coexpressed above random genes, as observed for the negative-loaded genes in excitatory neurons (third row).

#### **Supplementary Methods**

Correction of bulk tissue cellular composition effects

**Immunohistochemistry (IHC) analysis:** We obtained an estimate of cell type proportions for the ROSMAP samples from Patrick et al. (2020), who used immunohistochemistry (IHC) and microscopy to measure the proportions of up to five cell types: neuron, astrocyte, oligodendrocyte, microglia, and endothelial cells in 71 samples, of which 49 had measurements for all five cell types. We used the cell type proportions from Patrick et al. to test the impact of cellular composition variation (CCV) correction on coexpression. As in Chu et al. (2025), we performed correction by using a multivariate regression model to predict bulk level expression as a function of cell type proportions. Specifically, cell type proportions can be represented using six vectors, with each vector $\alpha_{k}$ containing the proportion of cell type $k$ across S subjects. Then, we modeled bulk tissue expression of gene A as follows:

$${Exp\_bulk}_{s}\left( A \right)=\beta_{0}+\sum_{k} \beta_{k}*\alpha_{k,s}+\epsilon_{A,s}$$

where $\alpha_{k,s}$ denotes the proportion of cell type k in subject s, $\beta's$ are the model coefficients, and $\epsilon_{A,s}$ denotes the residual error. This model allowed us to measure the proportion of variance, $R^{2}$, in the bulk expression of A that can be explained by CCV. The residuals of the regression models represent a removal the estimated effects of CCV. We compared coexpression before and after correction for CCV.

**Marker Gene Profile (MGP) analysis:** We selected genes as cell type markers for estimating the MGPs as follows, from the Velmeshev and ROSMAP single cell datasets. First, we computed the mean per-cell type expression levels for each gene within each dataset and selected the top 100 genes in each cell type based on the minimum fold change over other cell types. That is, the difference of expression between each cell type and the maximum expression among all other cell types. This approach prioritizes genes with cell type-specific expression, instead of merely those with the highest expressions in the given cell type. For each cell type, we selected the genes identified in both datasets as the final set of marker genes. The final set has 381 genes including 42 for excitatory neuron, 62 for inhibitory neuron, 66 for astrocyte, 71 for microglia, 71 for oligodendrocyte, and 69 for OPC. Using these marker genes, we estimated marker gene profiles using the method of Mancarci et al. (2017). Briefly, the expression of the marker genes for each cell type is summarized by PCA, and the first PC scores represent the MGPs for each cell type, interpreted directly as a proxy measure of relative cell type proportions.

#### Clustering and cluster integrity

To identify xCell clusters for each cell type, we first, derived a consensus for each cell type by aggregating the coexpression matrices of different datasets. We performed this aggregation by averaging the rank-transformed coexpression values across the given set of matrices followed by an additional round of rank normalization. After this procedure, there was a consensus for each cell type. Next, we converted coexpression values into distance measures by using the equation: $f(x) = (1 - x)$. We used WGCNA’s hierarchical clustering algorithm to identify coexpression clusters, via the dynamic tree cutting algorithm with a stringent distance cut-off threshold of 0.2 to pinpoint the strongest clusters. This parameter ensures that the average distance of the genes to the centroid of its cluster would be the distance defined by the cut-off threshold. Finally, we used the EGAD algorithm (Ballouz et al., 2017) to measure cluster integrity. This algorithm has been used in several previous studies to quantify the cluster integrity of genes annotated with the same functions (Crow et al., 2016; Ballouz et al., 2017; Skinnider et al., 2019; Harris et al., 2021). Because this method can be used to measure the integrity of any arbitrary cluster, we adopt it here to measure the preservation of the xCell coexpression clusters at the xSubject and xBulk levels. EGAD uses area under the receiver operating characteristic curve (AUROC) to measure the extent that the genes within a given cluster are connected with each other and disconnected with genes in other clusters.

#### Using xSubject expression to explain xBulk expression profiles.

To examine the effects of dilution at the xBulk level, we used the xSubject expression vectors to predict bulk tissue expression profiles. Specifically, for each gene, we used a multivariate linear regression model to explain xBulk expression profile as a function of the six cell-type specific xSubject expression vectors. The xSubject level expression profile of gene A can be represented as an expression matrix with K rows (cell types) and S columns (subjects) where each element ${Exp\_sbj}_{k,s}(A)$ is the expression of A in cell type k and subject s. Each row of this expression matrix is, then, a cell type specific xSubject expression vector of size S. We modeled the xBulk expression profile across S subjects as follows:

$${Exp\_bulk}_{s}\left( A \right)=\beta_{0}+\sum_{k} \beta_{k}*{Exp\_sbj}_{k,s} (A)+\epsilon_{A,s}$$

where ${Exp\_sbj}_{k,s} (A)$ denotes the expression of gene A in subject s and cell type k, $\beta's$ are the model coefficients, and $\epsilon_{A,s}$ denotes the residual error. Because this analysis requires all six cell types to have the same genes in each dataset, we reprocessed each data group to include the union of genes detected across all cell types. As an additional control we computed null distributions for each, using 100 random permutations where we shuffled the bulk tissue vectors randomly with respect to the xSubject vectors. To assess the explanatory power of single cell type specific xSubject expression profiles, we also computed a separate linear regression model that use each cell type specific xSubject expression profile as the single predictor. In this case the xBulk expression profile is modeled separately for each cell type k as follows:

$${Exp\_bulk}_{s}\left( A \right)=\beta_{0}+\beta_{k}* {Exp\_sbj}_{k,s} (A)+\epsilon_{A,s}$$

Together, we produced seven regression models for each gene: a full model that uses the complete set of all K cell type specific expression profiles and six cell type specific models that respectively use each of the six cell type specific xSubject expression profile as the predictor. Among the single cell type models for each gene, we selected the best performing single cell type model in terms of $R^{2}$for comparison with the full model. Because the full models use more covariates than the single cell type models, we used the Akaike information criterion (AIC) for comparison with the independent cell type models.

#### **Extended data and figures**

In this section we present details of our investigations into the relationships between bulk tissue coexpression (xBulk) and the xCell and xSubject modes. We consider the effects of dilution and cellular composition variability (CCV)-induced coexpression, and present further details on the specific patterns that are preserved in xBulk.

### Cellular composition variability-induced coexpression at the xBulk level drives concordance with xCell coexpression signals.

To study composition effects, we require estimates of cell type composition in each sample. First, we were able to make use of immunohistochemical (IHC) microscopy-based cell type proportion estimates for a subset (49 samples) of the ROSMAP cohort (Supplemenatry Methods) (Patrick et al., 2020). We used a multivariate linear regression model to quantify the degree to which the expression pattern of each of the 21,491 genes can be explained by the IHC cell type estimates (See Methods). On average, the IHC estimates could explain 16% of the total variance per gene (Extended Data Figure 1A), consistent with Patrick et al., with 4,186 (19.4%) of 21,491 genes significantly predictable by the IHC cell type proportion estimates at FDR < 0.1. This suggests that the impact of CCV was widespread, in agreement with our previous work (Farahbod and Pavlidis, 2020). In principle, genes that are more variable across cell types are more likely to be influenced by CCV. Accordingly, there was a correlation (Pearson’s r = 0.14) between cross-cell type variance and the proportion of variance explained by cell type proportion estimates (Extended Data Figure 1A).

In parallel, we used a second approach for cell type composition estimation that was applicable to both the ROSMAP and Velmeshev data sets. Specifically, we used the marker gene profile (MGP) method which makes estimates of relative proportions directly from the expression data (Mancarci et al., 2017). Its drawback, like most such “deconvolution” methods, is that it is less direct than physical counts of cells. In comparison to the IHC models, the MGPs, on average, could explained a higher proportion of variance per gene (mean $R^{2} = 0.36$) (Extended Data Figure 1B). In part due to the large sample size (636 samples), most genes (20,330 of 21,491) were significantly predictable at FDR < 0.1. We also found a stronger correlation (Pearson’s r = 0.38) between cross cell type variability and proportion of variance explained per gene (Extended Data Figure 1B). This was reproduced in the Velmeshev data as well (Extended Data Figure 2). Here, MGPs could explain, on average, $R^{2} = 0.41$ of the total variances per gene and 16,630 (of 25,326) genes were significantly predictable at FDR < 0.1 (Extended Data Figure 2).

Next, we attempted to reduce the effects of CCV in the 49 bulk samples by extracting the residuals of the multivariate linear models, producing a “corrected” version of the expression data (tested with both MGP and IHC estimates for ROSMAP and with MGPs for Velmeshev). Specifically, we looked at the change in the correlation values for the edges that were identified in the xCell and xSubject coexpression. In excitatory and inhibitory neurons the average coexpression values of xCell edges underwent clear reductions upon CCV correction (Extended Data Figure 1C). On the other hand, the xSubject edges for these cell types remained relatively intact. These observations were corroborated by the MGP analysis, in that xCell edges for excitatory and inhibitory neurons became less co-expressed after CCV correction (Extended Data Figure 1C), and the changes were larger than using IHC estimates. Despite being more aggressive, the impact of this procedure on xSubject edges remained minimal. The reduction in xCell edge co-expression also extended to other cell types with the exception of OPCs. Regressing out the MGPs in the Velmeshev dataset, again, led to reductions in co-expression of xCell edges, especially in excitatory and inhibitory neurons (Extended Data Figure 1C). Importantly, after correction using MGPs, xBulk in both datasets had higher concordance with xSubject than with xCell for all six cell types (Extended Data Figure 1D). This observation was consistent with the intermediate position of xSubject between xBulk and xCell levels.

These observations suggest that the apparent preservation of xCell edges were driven, at least in part, by CCV. Since CCV is a proxy for differential expression among cell types (Farahbod and Pavlidis, 2020), there may be a nontrivial relationship between cell-type-specific expression and intracellular regulation. In other words, genes specifically expressed in a given cell type may be more likely to be dynamically co-regulated within that cell type.

### Few xCell coexpression gene clusters were preserved at the xSubject and xBulk levels

The comparisons thus far ignore differences among genes. We hypothesized that there would be particular clusters of genes with higher coexpression signal preservation than others. We identified coexpression clusters at the xCell level to evaluate their preservation at the xSubject and xBulk levels as follows. For this analysis we could only consider the 7,868 genes detected in all datasets for each cell type. First, we produced a consensus for each cell type by aggregating xCell across datasets. Then, we performed hierarchical clustering to identify a set of coexpression clusters in each cell-type-specific consensus. We chose a stringent cut height of 0.2 (rank normalized coexpression) and a minimum cluster size of 30 to pinpoint the strongest coexpression clusters in each of the six cell types (see Methods for details). Across all cell types, we obtained 115 clusters. Note that the clusters identified in different cell types may contain some of the same genes, though such overlaps were limited. Overall, 5,164 of the 7,868 genes considered were members of at least one cluster. Within each cell type, between 16% (microglia) to 57% (excitatory neurons) of genes belonged to a cluster (low gene coverage was expected due to the stringent threshold). There were two large clusters containing 1,059 genes and 911 genes which were respectively identified in excitatory and inhibitory neurons. The remaining clusters contained 70.9 genes on average (min = 30; max = 248). Analysis of a subset of the clusters (those showing highest and lowest preservation at xBulk; see below) is shown in Extended Data Figure 3. Extended Data Figure 3A shows the sizes of these clusters.

As a first step to ensure the quality of the clusters at the xCell level, we used a guilt-by-association (GBA) approach with the EGAD algorithm (see Supplementary Methods). This summarizes both intra-cluster cohesion and inter-cluster separability for each cluster with an EGAD AUROC value. AUROCs range from range from 0 to 1 with values above 0.5 indicating higher performance than random chance. As expected, across all cell types and datasets, the average AUROC was above 0.9 for all cell types in the consensus where the clusters were identified. Further, the performance of each cluster within the corresponding cell type was robust across the components (Extended Data Figure 3B, top).

Next, we looked at the preservation of xCell clusters both at the xSubject and at xBulk levels using EGAD. Most clusters had AUROC above random chance (>0.5) at both xSubject and xBulk levels (Extended Data Figure 3B, middle and bottom). However, on average, AUROC decreased from 0.92 at the xCell level to 0.81 at the xSubject level and finally 0.72 at the xBulk level. Cluster inhibitory.id_0018 exemplified this pattern. This cluster of genes was highly and reproducibly co-expressed in xCell (mean AUROC = 0.92), but its integrity decreased considerably at the xSubject level (mean AUROC = 0.70). Finally, at the xBulk level, the genes within inhibitory.id_0018 had little discernible coexpression (mean AUROC = 0.52).

Despite the general reduction of cluster integrity, there were some clusters that were exceptionally well preserved across levels. This included clusters that contain ribosomal protein genes (RPGs) including microglia.id_0001, opc.id_0001, astrocyte.id_0002, and oligodendrocyte.id_0001 (Extended Data Figure 3A). RPGs are known to exhibit robust coexpression in xCell and xBulk (Li et al., 2016). Other examples were excitatory.id_0010 and excitatory.id_0020 which had AUROC of >0.9 in all of xCell, xSubject, and xBulk in both Velmeshev and ROSMAP datasets (Extended Data Figure 3B). Overall, there were five (of 115) clusters with AUROC of >0.9 at the xBulk level in at least one dataset, exemplifying exceptional preservation of a subset of xCell coexpression patterns.

### Correction of bulk tissue for cellular composition reduces cluster preservation

Above we showed that CCV can drive concordance of xBulk with xCell. To investigate this further, we looked at the change in the average intra-cluster coexpression values before and after statistical correcting for CCV effects. Here, we used ranked normalized coexpression values instead of EGAD AUROC in order to assess the changes of coexpression within each cluster in isolation from other genes. We performed three separate CCV correction experiments using the IHC data in ROSMAP and MGPs in both ROSMAP and Velmeshev datasets. We selected the top 20 clusters with the highest xBulk EGAD AUROCs for analysis. There were a number of clusters with substantial reductions in coexpression following CCV correction (Extended Data Figure 3C). Notably, the coexpression of genes in oligodendrocyte.id_0010 and oligodendrocyte.id_0014 decreased by 0.20 and 0.24 after MGP correction in Velmeshev, but not ROSMAP. This was consistent with the previous observation that CCV appeared to have a stronger influence on oligodendrocyte associated expression in the Velmeshev dataset (Extended Data Figure 1C and D). The genes in these clusters were also particularly highly expressed in oligodendrocytes (Extended Data Figure 3D). The most highly connected genes in oligodendrocyte.id_0014 were associated with myelination. Beyond the oligodendrocyte specific clusters, there were three other clusters that lost coexpression upon CCV correction: excitatory.id_0023, microglia.id_0006, and astrocyte.id_0007. The reduction of coexpression in these clusters were reproducible across all three analyses (Extended Data Figure 3C). The genes in these clusters were also particularly highly expressed in neurons, making them more susceptible to CCV influence (Extended Data Figure 3D).

The reduction in xBulk coexpression following removal of CCV suggest that the preservation of these instances of xCell coexpression was potentially driven primarily by different sources: one is due to cell-to-cell variability *within* a type, and the other is evident across subjects in bulk tissue due to cell-type specificity of the same genes. However, for these clusters, the coexpression at the xBulk level was not directly related to that observed at the xCell level. Consider the clusters astrocyte.id_0007 and microglia.id_0006, which were respectively identified at the xCell level in astrocyte and microglia. Despite glial cell-specific xCell coexpression, the expression of the genes within these clusters was particularly elevated in neurons, resulting in CCV-induced coexpression at the xBulk level. For example, the most highly coexpressed genes in astrocyte.id_0007 included genes associated with synaptic functions. Although it is possible that these clusters reflect contamination of ambient neuronal mRNA, they demonstrate the potential mismatch between xCell coexpression and cell type expression specificity. In other words, stable and specific expression of genes in a particular cell type does not necessarily indicate coexpression of those genes within the same cell type.

CCV could not explain all of the observed cluster preservation at the xBulk level. In particular, CCV corrections had small effects on the clusters that contained ribosomal protein genes: microglia.id_0001, opc.id_0001, astrocyte.id_0002, and oligodendrocyte.id_0001 (Extended Data Figure 3A and C). CCV correction also appeared to improve intra-cluster coexpression among genes in clusters excitatory.id_0020, excitatory.id_0010, and inhibitory.id_0012 (Extended Data Figure 3C). These findings suggest that their preservation originated from the xSubject level. These clusters represent a subset of coexpression patterns that may have actually propagated from the xCell level to the xSubject and xBulk levels.

### Cell type specific xSubject level expression patterns undergo dilution at the bulk level

Dilution of cell type specific signals, due to the presence of other cell types in the sample, obscures the propagation of cell-type-specific xSubject coexpression signals to the xBulk level. The availability of matched subjects in both single cell and bulk tissue offers an opportunity to directly explore the relationship of expression patterns between the xSubject and the xBulk levels. In Chu et al. (2025) we devised and demonstrated a model that relates xBulk expression to the underlying cell-type-specific xSubject expressions, which we reapply here. Essentially, we proposed that the bulk level expression of a given gene in a given sample is the weighted sum of the cell type specific subject level expressions of the gene, where the weights are the cell type proportions. Under this model, xBulk expression patterns are strictly the result of aggregating cell type specific xSubject expression vectors. Accordingly, we refer to it as the “dilution model” because the xSubject variation of any one cell type is expected to undergo dilution upon aggregation with other cell types. This model ignores the contribution of CCV, which we have already shown to be an important source of variability.

Using this model, we compared the xSubject and xBulk expression patterns in both ROSMAP and Velmeshev datasets. For each gene, we used a multivariate linear regression model to explain xBulk expression based on the combination of six cell type specific xSubject expression vectors (See Methods). If the dilution model is at least partly correct, the xSubject expression vectors would be a significant source of variability at the xBulk level. The model could explain 26.6% and 24.8% of the total variance in the ROSMAP and Velmeshev datasets respectfully across all the genes analyzed (Extended Data Figure 4A and B). There was a clear rightward skew in the p-value distributions in both datasets (Extended Data Figure 4C and D), though statistical significance was more apparent in the Velmeshev dataset, where 1,679 (12.4%) genes were statistically significant at FDR < 0.1, than in the ROSMAP dataset where only 29 (0.29%) were significant. As a control, we shuffled the xBulk expression vectors randomly (100 times) to compute the null distribution for each gene. Reassuringly, the null p-value distribution was nearly uniform across all 100 iterations in both datasets (Extended Data Figure 4C and D). Further, the null models could explain, on average, only 16.7% and 22.9% of the total variance in the Velmeshev and the ROSMAP datasets, respectively. Finally, 1,311 (9.67%) genes in the Velmeshev dataset and 249 (2.50%) genes in the ROSMAP dataset had higher] $R^{2}$ values than all 100 $R^{2}$ values generated using the shuffled data. Taken together, the xBulk expression patterns could be significantly explained by the xSubject expression vectors, corroborating the dilution model.

Given the cell type heterogeneity of brain cortical tissue, it is likely that no one cell type would dominate the expression pattern at the xBulk level. To explore this, we looked at the proportion of variance that could be explained independently by the xSubject expression profile of each cell type. The independent models use a cell-type-specific xSubject expression vector as a single predictor to explain xBulk expression. On average, excitatory neuron was the most explanatory cell type in both datasets. In the Velmeshev dataset, excitatory neuron was the maximally explanatory cell type in 3,928 (29%) of all genes in the Velmeshev dataset where it could explain, on average, $R^{2} = 0.18$ of the total variances. In contrast, OPC was the maximally explanatory cell type in only 1,529 (11.3%) genes and could only explain $R^{2} = 0.11$ of the total variances. We found comparable numbers in the ROSMAP dataset as well. This was expected given the relatively larger cell type proportion of excitatory neurons. Nonetheless, the maximally explanatory cell type clearly varied across genes (Extended Data Figure 4E and F). Importantly, for many genes, the full models were substantially more explanatory than the single covariate models that uses the best performing cell type (Extended Data Figure 4E and F). This was consistent with the dilution model where the xBulk level contains contributions from multiple cell types. However, the full models may fit the data more closely due to increased complexity since each full model uses six predictors. To account for this we used Akaike Information Criterion (AIC) to compare the full models with the best performing single covariate counterparts for each gene. The majority of genes (Velmeshev: 89.1%; ROSMAP: 93.2%) had lower AICs in the single covariate model than the full model (Extended Data Figure 4G and H). There were just 75 genes in the Velmeshev dataset and 19 genes in the ROSMAP dataset where the AIC of the full model was smaller than the best performing single model by 10 units, validating the dilution model. In other words, the xBulk expression patterns for these genes could not be traced back to any particular cell type. Rather, they were a combined version of several cell type specific xSubject expression patterns, lending further support to the dilution model.

Despite the effects of dilution, a key prediction of the dilution model is that for genes with highly specific expression in a particular cell type, its expression pattern in the given cell type would dominate variability at the xBulk level. Consistent with this hypothesis, genes maximally explained by each cell type had elevated levels of expression within the same cell type (Extended Data Figure 4I and J). However, cell type expression specificity cannot explain the exceptional preservation of coexpression clusters that could not be attributed to CCV, because the expressions of these genes were not particularly elevated in any cell type (Extended Data Figure 3D). These observations suggest these coexpressions may have been maintained due to inter-cell type synchrony (Chu et al. 2025). To test this hypothesis, we looked at the correlation of xSubject expression profiles among the different cell types (See Methods). Remarkably, the aforementioned clusters were highly synchronized among the different cell types (Extended Data Figure 3E). We additionally confirmed the presence of inter-cell type synchrony for these clusters in all seven datasets. The presence of inter-cell type synchrony provided an explanation for the preservation of coexpression signals of these xCell clusters at the xBulk level.

To investigate whether inter-cell type synchrony can broadly explain preservation of xSubject coexpression signals at the xBulk level, we looked at the association between inter-cell type synchrony and preservation of coexpression signals. First, we separated the genes into ten equal size bins (Velmeshev: 1,377 genes; ROSMAP: 996 genes) with different levels of inter-cell type synchrony. The top bin contained genes with biological sex-specific expression profiles. Because these genes were differentially expressed between sexes in every cell type, they exemplified an extreme version of inter-cell type synchrony. Importantly, the majority of RPGs in both datasets were also found in the top bin, hence providing an explanation for their preservation of coexpression patterns (Extended Data Figure 5). Next, we directly measured the correlation of coexpression between xSubject and xBulk for the genes in each bin. We found a clear relationship between inter-cell type synchrony and coexpression concordance between xSubject and xBulk coexpressions (Extended Data Figures 6 and 7). In other words, xSubject coexpression relationships remained largely intact at the xBulk level for genes with high levels of inter-cell type synchrony, which we interpret as allowing them to escape the effects of dilution.

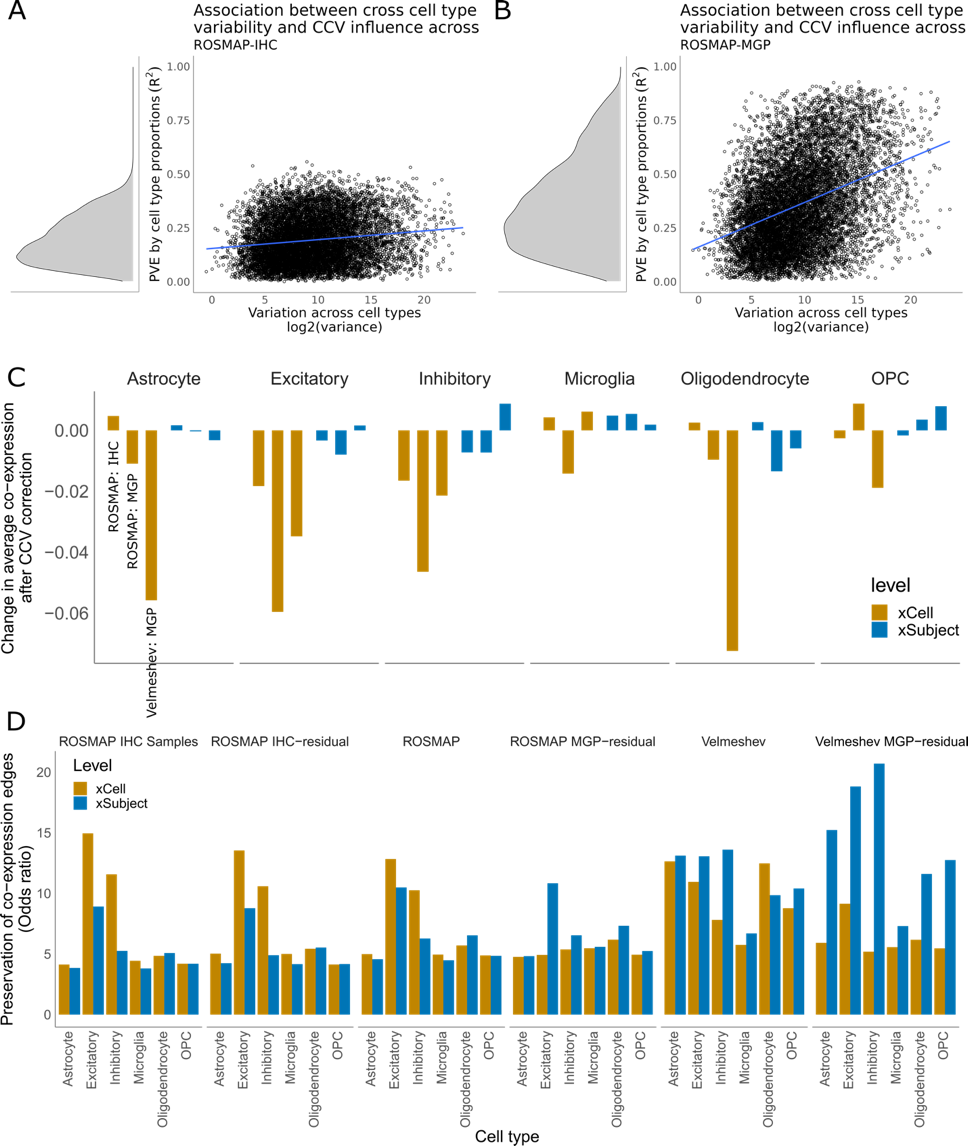

**Extended Data Figure 1. Correction for composition variation reduces preservation of xCell signals at the xBulk levels:** (A) Scatterplot showing the relationship between variability of mean expressions across cell types (x-axis) vs. proportion of variance that could be explained by the models that use IHC-estimated cell type proportions (y-axis) in ROSMAP. Each datapoint is a gene. In general, the more variable a gene is across cell types, the more likely its xBulk expression would be driven by CCV. (B) Same as (A) but for models that use MGPs instead of the IHC-based estimates. The same plot for Velmeshev is shown in Extended Data Figure 2. (C) Bar plots showing the change in correlation of expressions of edges that were identified in xCell and xSubject before and after CCV correction. Separate bars shown from different corrective methods and datasets. A large negative change indicates that CCV is an important driver for coexpression of the given edges at the xBulk level. (D) Bar plots showing odds ratios measuring preservation of xCell or xSubject coexpression at the xBulk level in both ROSMAP and Velmeshev datasets, before and after CCV correction. Results for ROSMAP IHC samples are plotted in a separate panel because IHC-based cell type proportion estimates were only available for 49 of all 636 samples in ROSMAP.

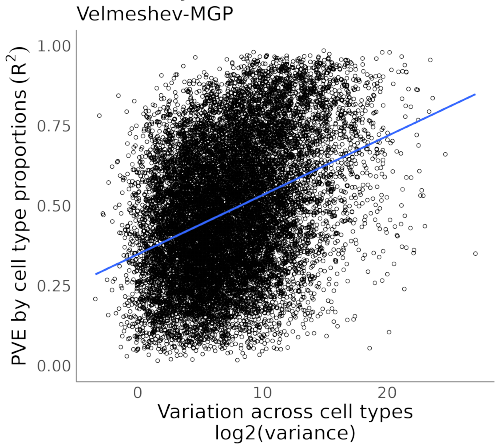

**Extended Data Figure 2: Relationship between gene variation across cell types and MGP-estimated CCV effects for the Velmeshev data set**. As for Extended Data Figure 1B, but for the Velmeshev data.

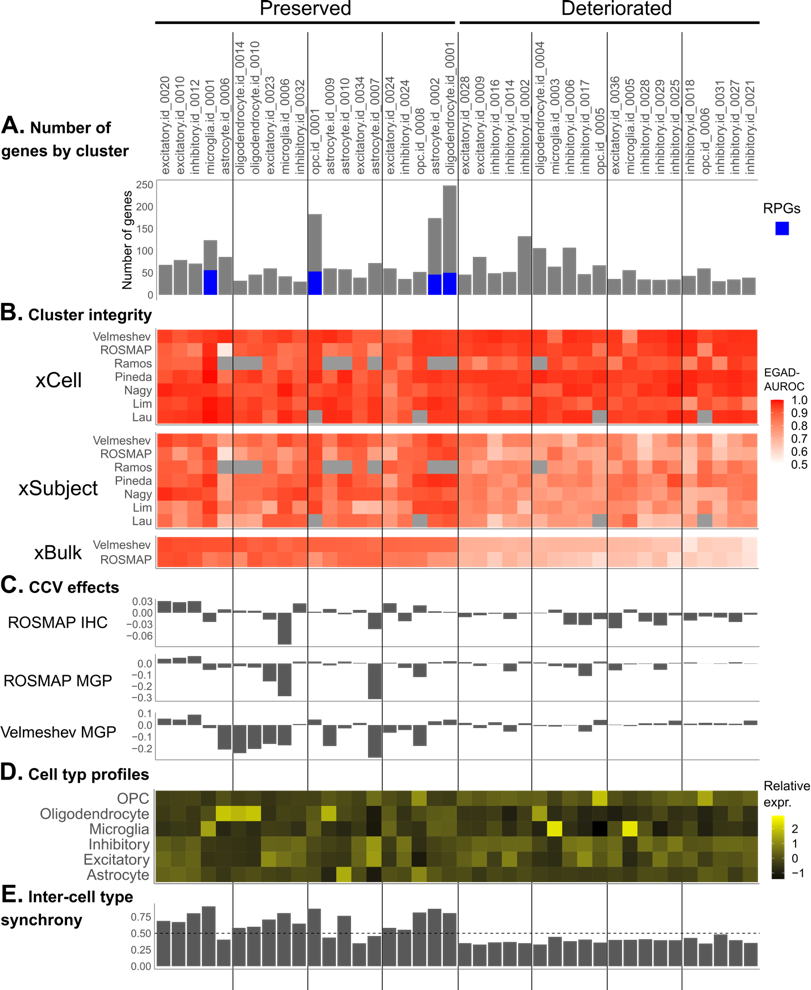

**Extended Data Figure 3 Preservation of some clusters is driven by CCV effects:** Cluster properties for the 20 most highly preserved (left) and 20 least preserved (right) xCell coexpression clusters at the xBulk level. (A) Bar plots showing the number of genes in each cluster. The number of ribosomal protein genes (RPGs) is shown in blue. (B) Heatmap showing the cluster integrity, measured by EGAD AUROC, of each cluster in each dataset at the three levels of analysis. For the xCell and xSubject levels, AUROC is computed only for the cell type of origin for the given cluster. At the xCell level, all clusters had high AUROCs as expected. A number of clusters lose their coexpression at the xSubject and xBulk levels. Missing values are indicated by grey squares. (C) Bar plots showing the change in average intra-cluster correlations at the xBulk level before and after CCV correction based on IHC or MGPs and different datasets. (D) Heatmap showing the mean expression of genes in each cluster across different cell types in Velmeshev, each column is centered to 0. (E) Bar plots showing the level of average inter-cell type synchrony (rank normalized correlation, 0.5 is expected at random) of the genes in each cluster in Velmeshev. Many of the highly preserved clusters contain genes with elevated levels of inter-cell type synchrony at the xSubject level.

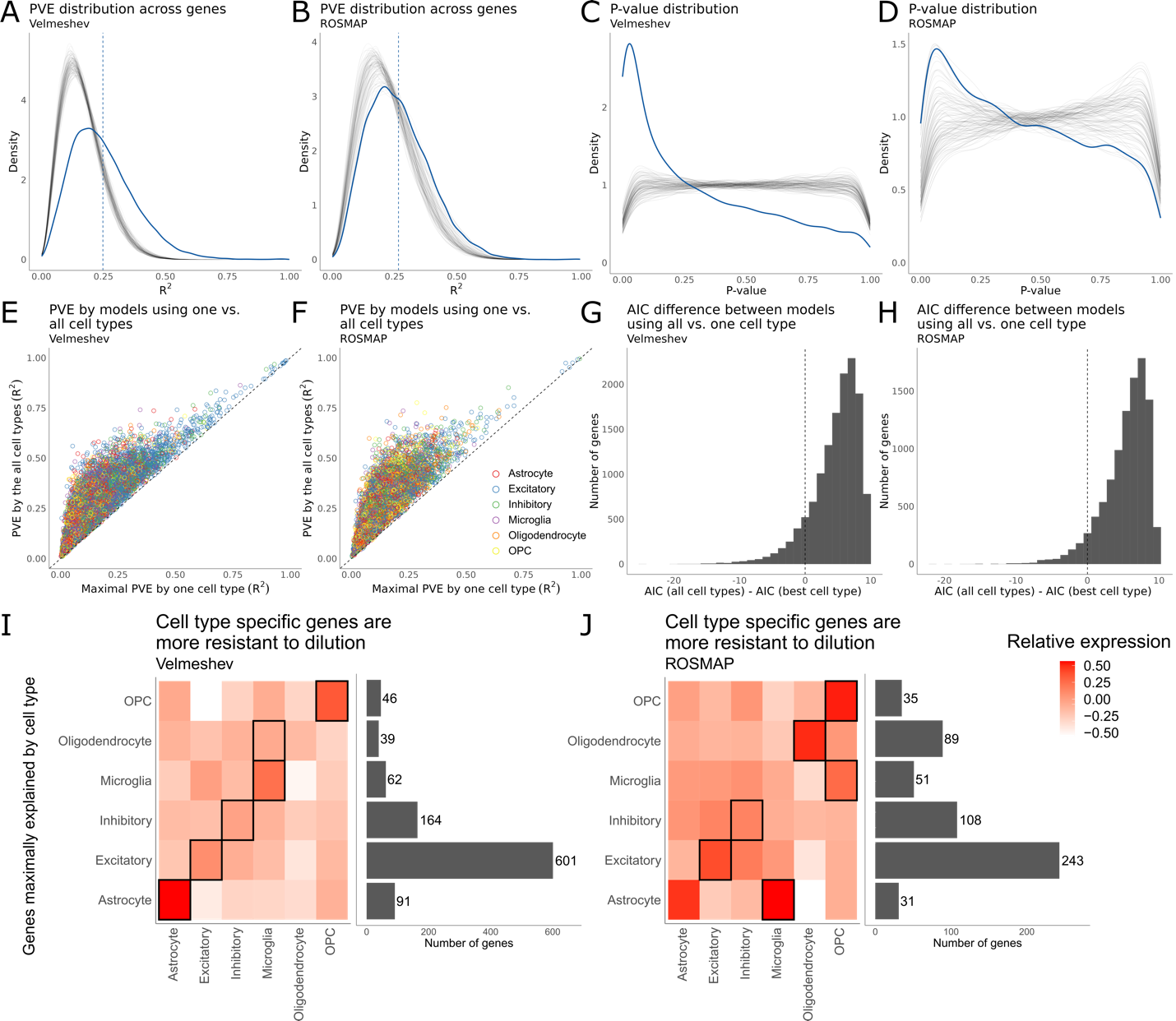

**Extended Data Figure 4: The effects of dilution of cell type specific xSubject expression patterns** (A-B) Distribution of proportion of variance per gene that could be explained by the dilution models using xSubject expressions as predictors. The blue line plots the distribution of the observed values across genes. The grey lines are the 100 null distributions generated upon randomly shuffling the subjects. (C-D) Distribution of the corresponding model p-values, for the same models plotted in (A-B). (E-F) Scatterplot comparing Proportion of variance explained (PVE) of the best performing independent cell type models (x-axis) against the full models (y-axis) that use expressions of all the cell types. (G-H) Distributions of difference of AIC between the full model and the best performing independent model. (I-J) Heatmap showing the cell type expression profile of genes maximally explained by each cell type at R^2^ > 0.3. The bars on the right show the number of genes maximally explained by each respective cell type on the y-axis. Each box shows the mean expression of those genes in each cell type (x-axis).

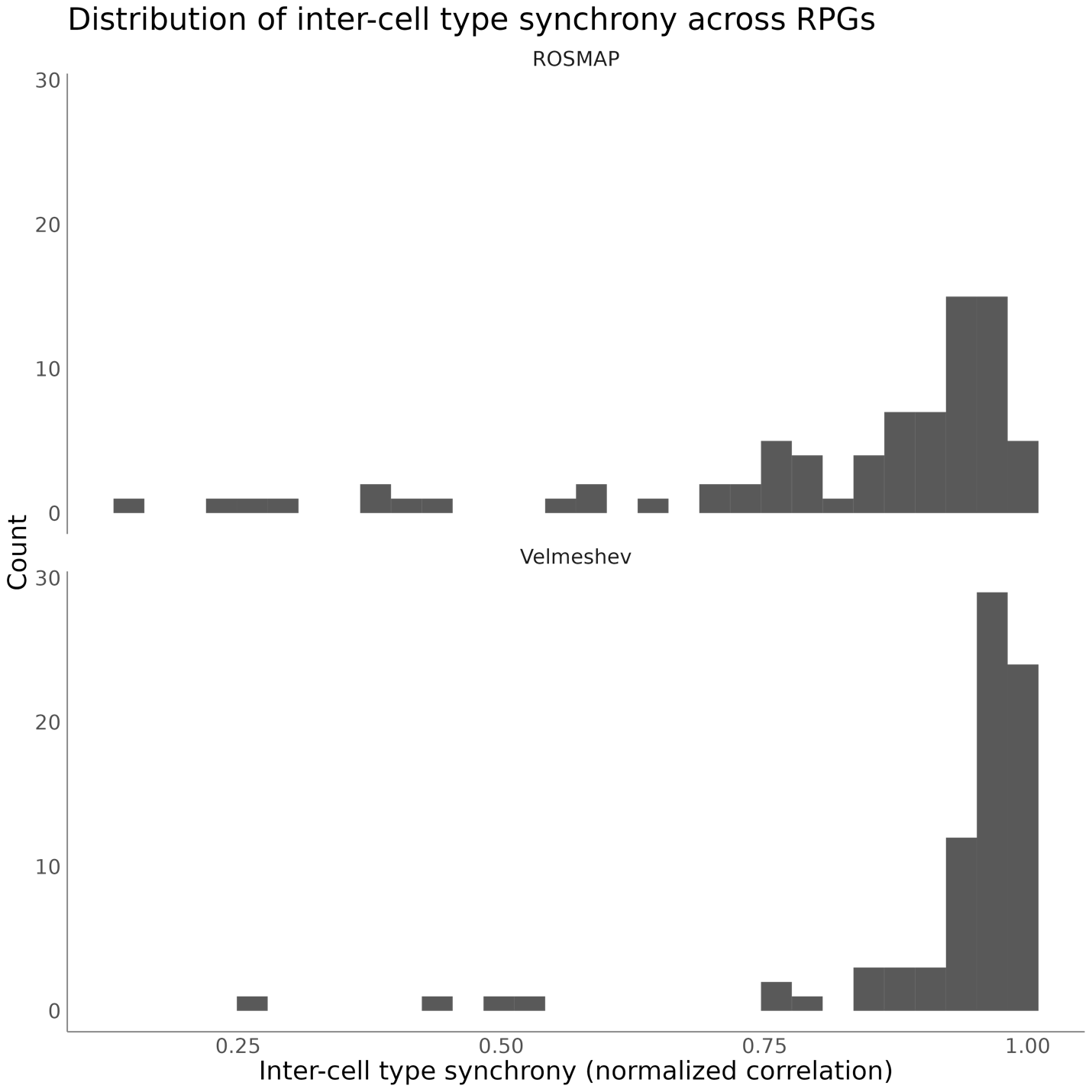

**Extended Data Figure 5: Inter-cell type synchrony of ribosomal protein genes (RPGs).** Histogram of distribution of normalized correlation of expression between cell types for RPGs. RPGs show high levels of inter-cell type synchrony in both ROSMAP (top) and Velmeshev (bottom), providing an explanation for their exceptional signal preservation from xCell to xBulk.

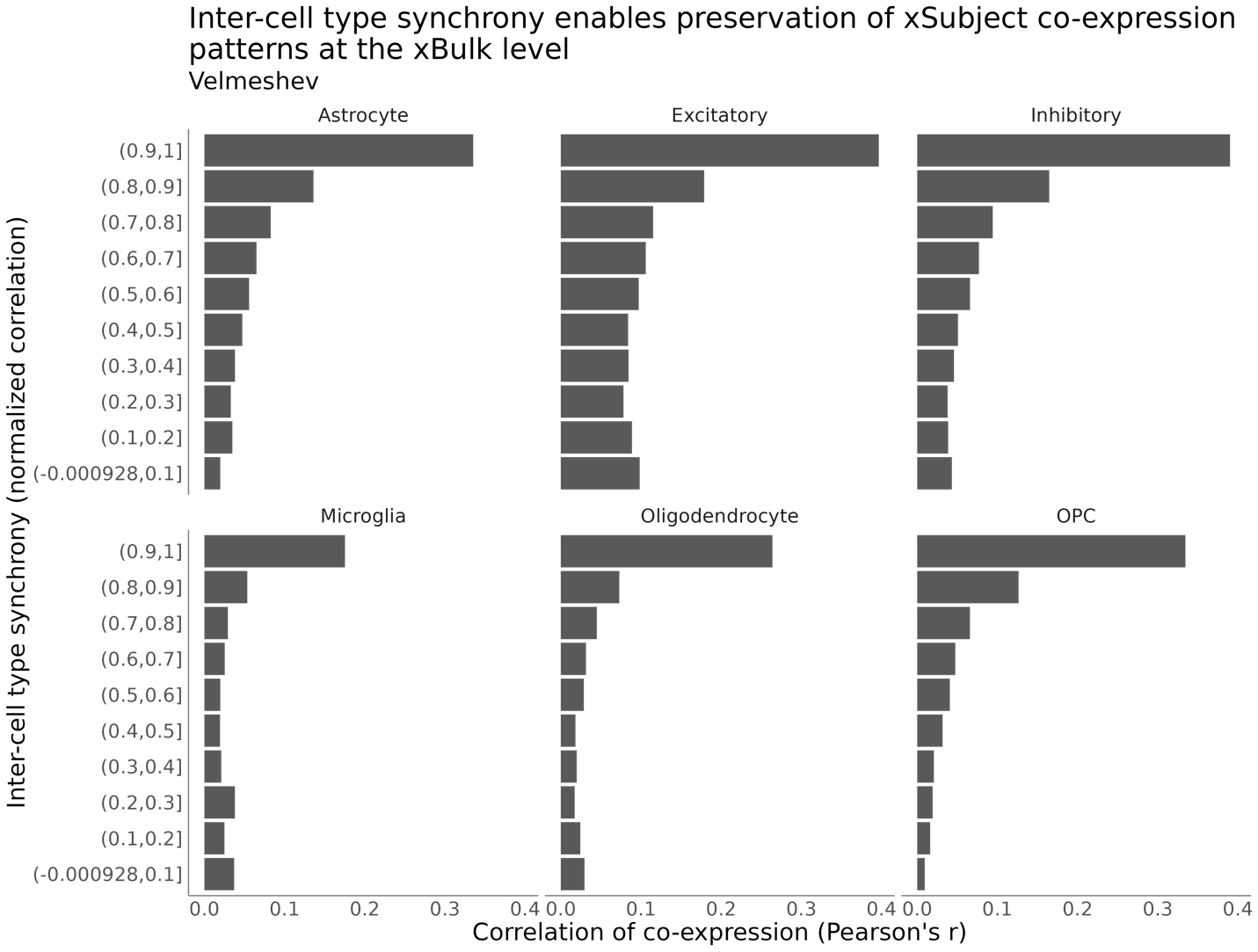

**Extended Data Figure 6: Association between inter-cell type synchrony and preservation of xSubject co-expression patterns in xBulk (Velmeshev).** Each bar shows “concordance of co-expression” or the level of correlation of co-expression values (x-axis) across gene pairs between the xSubject and xBulk levels for each decile of inter-cell type synchrony (y-axis). The higher the level of inter-cell type synchrony, the more co-expression signal is propagated from the xSubject to the xBulk levels.

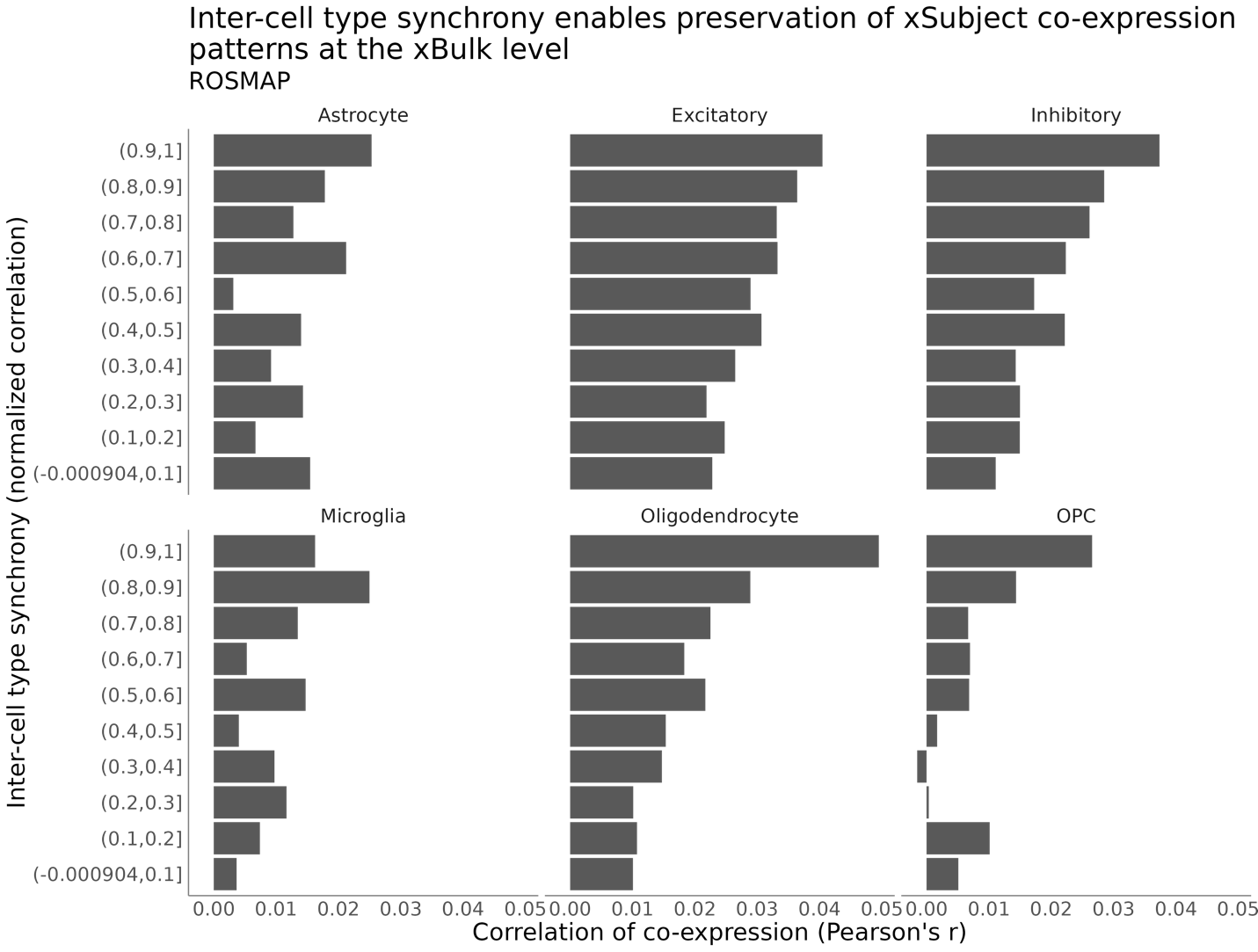

**Extended Data Figure Figure 7: Association between inter-cell type synchrony and preservation of xSubject co-expression patterns in xBulk networks (ROSMAP).** Same as Extended Data Figure 6 but for ROSMAP.

### **Supplement references**

Chu CP, Morin A, Pavlidis P (2025) Limits to the inference of gene regulation from bulk tissue expression data. :2024.10.24.619521 Available at: https://www.biorxiv.org/content/10.1101/2024.10.24.619521v3 [Accessed October 2, 2025].

Farahbod M, Pavlidis P (2020) Untangling the effects of cellular composition on coexpression analysis. Genome Res 30:gr.256735.119 Available at: http://genome.cshlp.org/lookup/doi/10.1101/gr.256735.119 [Accessed July 5, 2020].

Lau S-F, Cao H, Fu AKY, Ip NY (2020) Single-nucleus transcriptome analysis reveals dysregulation of angiogenic endothelial cells and neuroprotective glia in Alzheimer’s disease. PNAS 117:25800–25809 Available at: https://www.pnas.org/content/117/41/25800 [Accessed October 13, 2020].

Li X, Zheng Y, Hu H, Li X (2016) Integrative analyses shed new light on human ribosomal protein gene regulation. Sci Rep 6 Available at: https://www.ncbi.nlm.nih.gov/pmc/articles/PMC4921865/ [Accessed December 25, 2018].

Lim RG et al. (2022) Huntington disease oligodendrocyte maturation deficits revealed by single-nucleus RNAseq are rescued by thiamine-biotin supplementation. Nat Commun 13:7791 Available at: https://www.nature.com/articles/s41467-022-35388-x [Accessed August 7, 2024].

Mancarci BO, Toker L, Tripathy SJ, Li B, Rocco B, Sibille E, Pavlidis P (2017) Cross-Laboratory Analysis of Brain Cell Type Transcriptomes with Applications to Interpretation of Bulk Tissue Data. eNeuro:ENEURO.0212-17.2017 Available at: http://www.eneuro.org/content/early/2017/11/20/ENEURO.0212-17.2017 [Accessed November 20, 2017].

Mathys H, Davila-Velderrain J, Peng Z, Gao F, Mohammadi S, Young JZ, Menon M, He L, Abdurrob F, Jiang X, Martorell AJ, Ransohoff RM, Hafler BP, Bennett DA, Kellis M, Tsai L-H (2019) Single-cell transcriptomic analysis of Alzheimer’s disease. Nature 570:332–337 Available at: https://www.nature.com/articles/s41586-019-1195-2 [Accessed February 11, 2020].

Nagy C, Maitra M, Tanti A, Suderman M, Théroux J-F, Davoli MA, Perlman K, Yerko V, Wang YC, Tripathy SJ, Pavlidis P, Mechawar N, Ragoussis J, Turecki G (2020) Single-nucleus transcriptomics of the prefrontal cortex in major depressive disorder implicates oligodendrocyte precursor cells and excitatory neurons. Nature Neuroscience:1–11 Available at: https://www.nature.com/articles/s41593-020-0621-y [Accessed April 27, 2020].

Patrick E, Taga M, Ergun A, Ng B, Casazza W, Cimpean M, Yung C, Schneider JA, Bennett DA, Gaiteri C, De Jager PL, Bradshaw EM, Mostafavi S (2020) Deconvolving the contributions of cell-type heterogeneity on cortical gene expression Nie Q, ed. PLoS Comput Biol 16:e1008120 Available at: https://dx.plos.org/10.1371/journal.pcbi.1008120 [Accessed March 23, 2023].

Pineda SS et al. (2024) Single-cell dissection of the human motor and prefrontal cortices in ALS and FTLD. Cell 187:1971-1989.e16 Available at: https://www.cell.com/cell/abstract/S0092-8674(24)00234-4 [Accessed February 11, 2026].

Ramos SI, Mussa ZM, Falk EN, Pai B, Giotti B, Allette K, Cai P, Dekio F, Sebra R, Beaumont KG, Tsankov AM, Tsankova NM (2022) An atlas of late prenatal human neurodevelopment resolved by single-nucleus transcriptomics. Nat Commun 13:7671 Available at: https://www.nature.com/articles/s41467-022-34975-2 [Accessed May 21, 2024].

Velmeshev D, Schirmer L, Jung D, Haeussler M, Perez Y, Mayer S, Bhaduri A, Goyal N, Rowitch DH, Kriegstein AR (2019) Single-cell genomics identifies cell type–specific molecular changes in autism. Science 364:685–689 Available at: https://science.sciencemag.org/content/364/6441/685 [Accessed May 16, 2019].
